# Polymorphic ribonucleoprotein folding as a basis for translational regulation

**DOI:** 10.1101/2022.02.11.480102

**Authors:** Tom Dendooven, Elisabeth Sonnleitner, Udo Bläsi, Ben F. Luisi

**Affiliations:** Department of Biochemistry, University of Cambridge, Tennis Court Road, Cambridge CB2 1GA, U.K.

**Keywords:** co-transcriptional RNA folding, Crc, RNA chaperone Hfq, metabolic regulation, translational regulation, ribonucleoprotein assembly

## Abstract

The widely occurring bacterial RNA chaperone Hfq is a key factor in the post-transcriptional control of hundreds of genes in *Pseudomonas aeruginosa*. How this broadly acting protein can contribute to the regulation requirements of so many different genes remains puzzling. Here, we describe the structures of higher-order assemblies formed on control regions of different *P. aeruginosa* target mRNAs by Hfq and its partner protein Crc. Our results show that these assemblies have mRNA-specific quaternary architectures resulting from the combination of multivalent protein-protein interfaces and recognition of patterns in the RNA sequence. The structural polymorphism of the ribonucleoprotein assemblies enables selective translational repression of many different target mRNAs. This system suggests how highly complex regulatory pathways can evolve and be rewired with a simple economy of proteinogenic components.

**Graphical Abstract:** 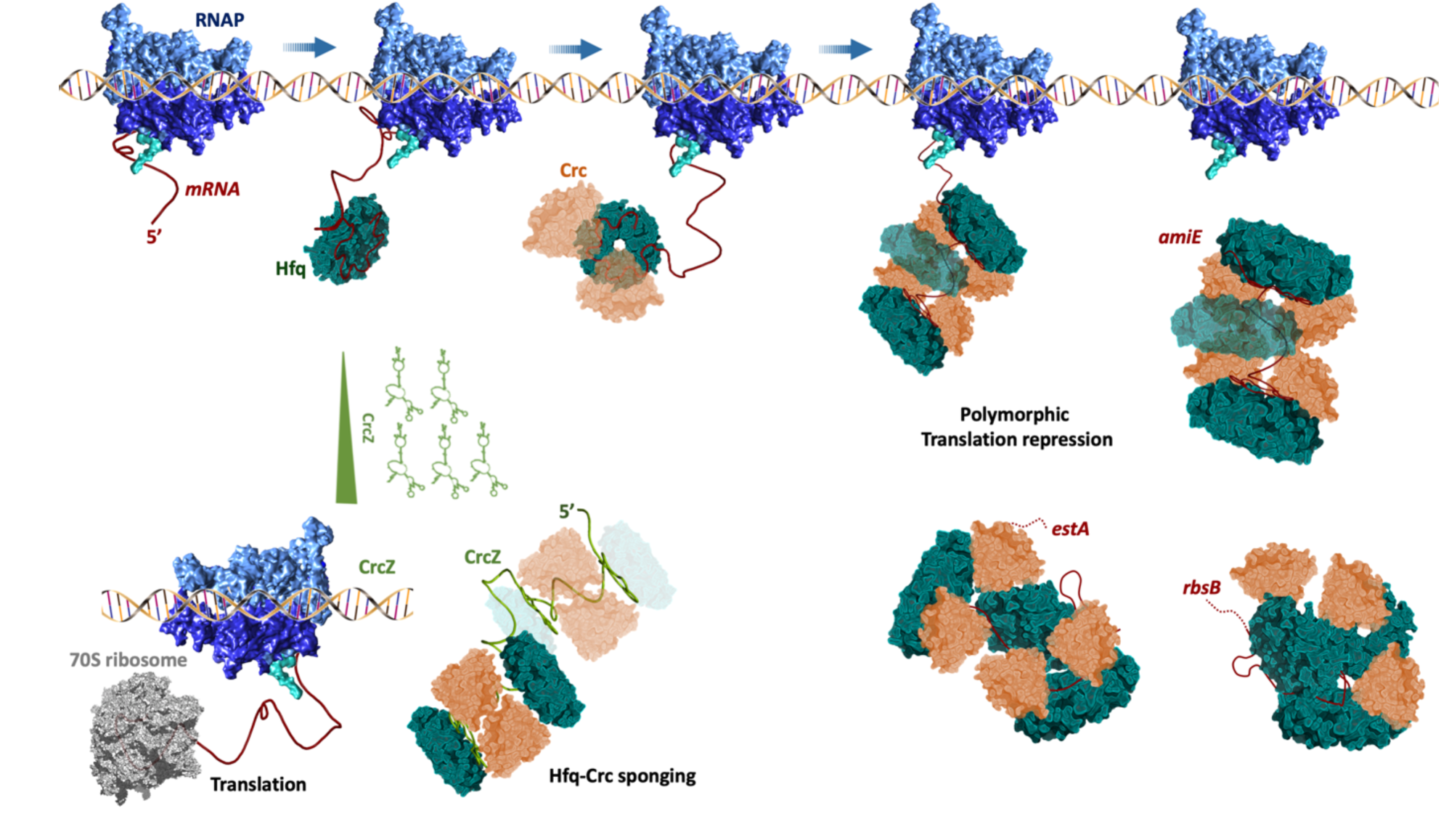

The RNA chaperone Hfq, in conjunction with the co-repressor Crc, forms higher order assemblies on nascent mRNAs. These complexes impact on translation of hundreds of transcripts in the pathogen *Pseudomonas aeruginosa*. Assemblies with different quaternary structures result from the interactions of the proteins with sequence motifs and structural elements in different mRNA targets, as well as from a repertoire of protein-to-protein interfaces. In this way, the combination of RNA sequence and two proteins can generate the diversity required to regulate many genes. It is proposed that the multi-step assembly process is highly cooperative and most likely competes kinetically with translation initiation to silence the targeted transcripts.

## Introduction

Two major post-transcriptional systems control gene expression in the pathogenic bacterium *Pseudomonas aeruginosa*. One system depends on the CsrA-like Rsm proteins, which act as translational repressors and engage GGA motifs in the Shine-Dalgarno sequence of target mRNAs that are often exposed in loops of stem-loop structures (Dubey *et al*., 2005; Schubert *et al*., 2007, Goodman *et al*., 2016; Holmqvist *et al*., 2016; Romero *et al*., 2018; Gebhardt *et al*., 2020). The other system depends on the RNA chaperone Hfq, a member of the widely occurring Lsm/Sm protein family, which facilitates the actions of small regulatory RNAs (sRNAs; Pusic *et al*., 2021), and can act as a translational repressor of target mRNAs (Sonnleitner & Bläsi, 2014; Sonnleitner *et al*., 2018; Kambara *et al*., 2018; Gebhardt *et al*., 2020; Malecka *et al*., 2021). Through these activities, Hfq contributes to the coordination of stress responses (Lu et al., 2016), metabolism (Sonnleitner & Bläsi, 2014), quorum sensing (Sonnleitner et al., 2006; Yang et al., 2015), virulence (Sonnleitner *et al*., 2003), and affects complex processes such as biofilm formation and the antibiotic susceptibility (Fernàndez *et al*., 2016; Pusic *et al*., 2016, Zhang *et al*. 2017; Pusic *et al*., 2018; Sonnleitner *et al*., 2020).

Hfq-mediated translational repression forms the basis for a hierarchical control of carbon and nitrogen utilization by *Pseudomonas spp*., a mechanism referred to as *c*arbon *c*atabolite *r*epression (CCR) (Rojo, 2010; Sonnleitner & Bläsi, 2014). CCR ensures that alternative nutrients are not utilized until the preferred carbon source, succinate, is depleted. The regulation is exerted through translational repression of genes affecting the uptake and metabolism of non-preferred nutrients (Sonnleitner & Bläsi, 2014). The best studied CCR-regulated gene is *amiE*, which encodes the enzyme aliphatic amidase that generates organic acids from short-chain aliphatic amides, thereby enabling *Pseudomonas* to utilize acetamide as a source of both carbon and nitrogen. When preferred carbon sources such as succinate are abundant, translation of *amiE* mRNA is suppressed through sequestration of the ribosome binding site by Hfq and the catabolite control protein Crc (Figure 1), which is followed by mRNA degradation (Sonnleitner & Bläsi, 2014). When the preferred carbon source is exhausted, CCR is alleviated by the regulatory sRNA CrcZ (Figure 1), which sequesters Hfq away from substrate mRNAs (Sonnleitner & Bläsi, 2014). The CrcZ levels are controlled by the alternative sigma factor RpoN (Sonnleitner *et al*., 2009; Abdou *et al*., 2011; Valentini *et al*., 2014) and the two-component system CbrA/B, which may be activated in response to the cellular energy status (Valentini *et al*., 2014).

**Figure 1:**
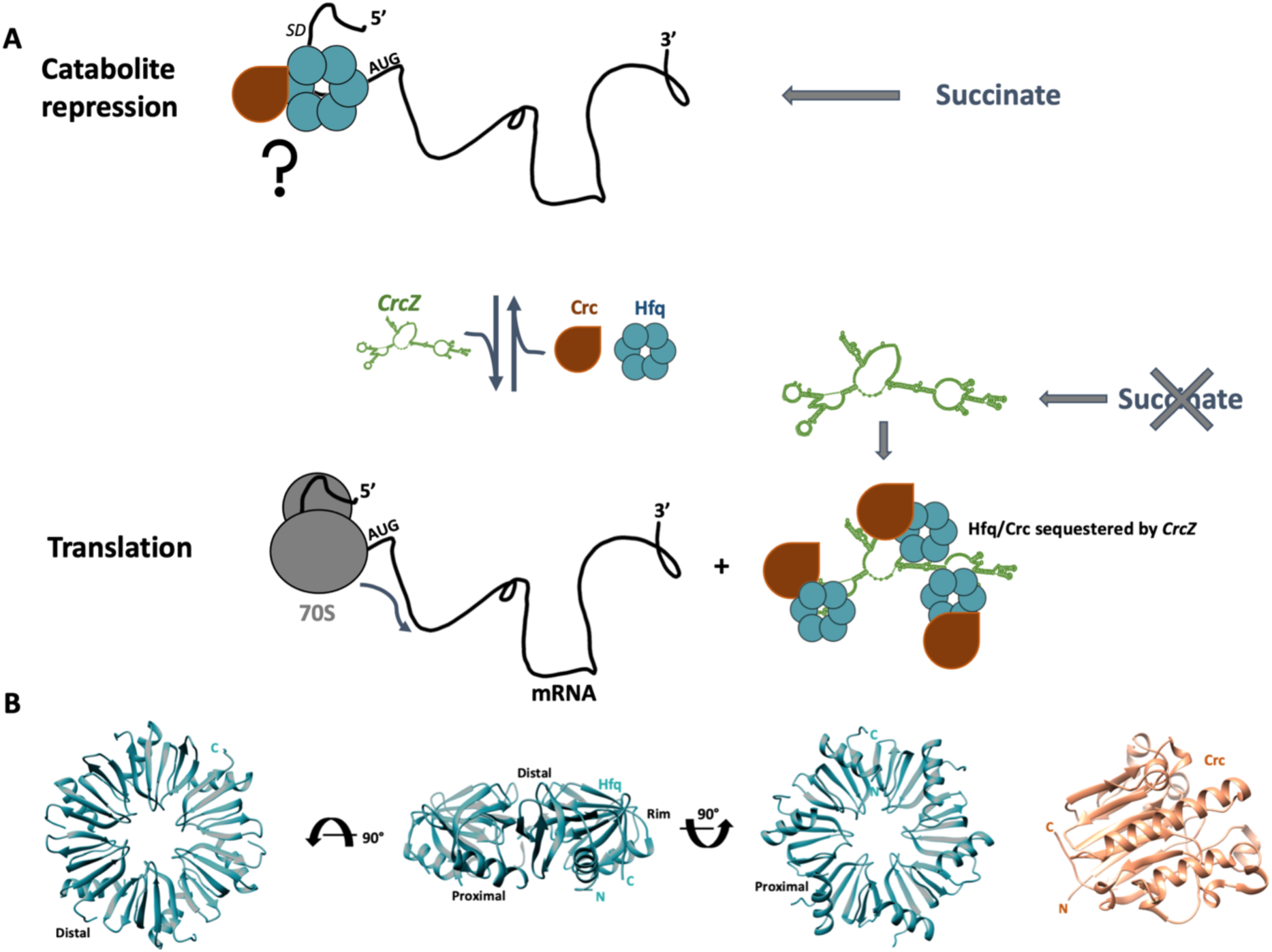
Catabolite repression in Pseudomonas spp. **(A)** During catabolite repression, e.g. when succinate levels are high (top), the Hfq hexamer and Crc cooperatively bind to A-rich sequences at the 5’-end of target mRNAs, and mask the ribosome binding site. When succinate levels are low (bottom), catabolite repression is alleviated by the regulatory RNA CrcZ, which sequesters Hfq and/or Hfq/Crc complexes. **(B)** Crystal structures of Hfq hexamer, showing the distal, proximal and rim side (left three panels), and Crc (right panel).

Understanding the molecular basis of CCR has been advanced by structural insights into RNA recognition by Hfq. These have identified three different RNA binding surfaces on the Hfq hexamer: the proximal face, the distal face, and the circumferential rim (Figure 1B). The proximal face binds uridine tracts, which are enriched at the 3’ end of sRNAs, the distal face has sequence preference for ARN triplet motifs (where A is an adenosine, R a purine and N any base), and the rim has arginine-rich patches that can interact with UA rich motifs of RNAs (Santiago-Frangos & Woodson, 2018). Cryo-EM structures of Hfq-Crc complexes on a short octadecameric segment derived from the 5’ upstream untranslated region (5’-UTR) of *amiE* mRNA (Supplementary Figure 1A; *amiE*_*6ARN*_) revealed how the Hfq distal side presents the *amiE* ribosome binding site to Crc (Pei *et al*., 2019). In these structures two stacked Hfq hexamers each present one *amiE*_*6ARN*_ motif to four Crc protomers. The structures suggested that translation repression complexes are higher order assemblies where several Hfq hexamers and Crc molecules are engaged on the mRNA target. How the Pseudomonad-specific Crc molecule contributes to the repressive function of these CCR assemblies has been a long-standing question. Interestingly, Crc has no intrinsic RNA binding capacity and does not interact with Hfq in the absence of a RNA substrate (Milojevic *et al*., 2013; Sonnleitner *et al*., 2018). Thus, the ability of Crc to engage Hfq-mRNA intermediates arises through the cooperative effects of the interactions in the higher order assemblies. In line with this model, single-molecule fluorescence assays and molecular dynamics simulations showed that Crc interacts with transient, pre-organized Hfq/RNA complexes and shifts the equilibrium towards assemblies with increased stability so that the effector complex blocks translation more efficiently (Malecka *et al*., 2021; Krepl *et al*., 2021). Thus, the cooperation of Hfq with Crc effectively stabilizes the repressive complex, excluding the 30S ribosomal subunit more effectively than Hfq alone.

Recent RNA-seq and proteomics approaches have revealed how CCR controls more than 100 mRNA targets that are co-regulated by Hfq and Crc, many of which are involved in carbon metabolism and virulence (Kambara *et al*., 2018; Corona *et al*., 2018). However, the diverse regulatory functions of Hfq/Crc present a puzzling issue: how can the same two effector molecules target so many different mRNA sequences with specificity and individually tuned response? To explore this question, we solved the cryo-EM structures of Hfq/Crc translation repression complexes assembled on extended 5’-end segments of the *amiE, estA* and *rbsB* genes, encoding an amidase, an esterase and a putative ribose transporter (Winsor *et al*., 2016), respectively, and present *in vivo* observations supporting our models. These structures revealed how multiple ARN repeats in the RNA targets are engaged by Hfq and Crc to form higher order repressive complexes. Strikingly, these complexes are polymorphic in quaternary organization. The determinants for the organization are a combination of RNA secondary structure elements and the position and length of the ARN motifs in the *t*ranslation *i*nitiation *r*egion (TIR) of these transcripts. Permissive Crc dimerization- and small, versatile Hfq-RNA-Hfq interfaces further stabilize the diverse repressive assemblies. The polymorphic character of these complexes enables Hfq and Crc to target many genes, while maintaining sequence specificity. These findings define a new paradigm for *in vivo* action of Hfq through cooperation with the Crc helper protein to form diverse RNA-driven effector assemblies that regulate expression of numerous target genes.

## Results

### Higher order Hfq/Crc assemblies form on mRNA targets *in vitro*

Based on our earlier cryo-EM structures of a complex formed by Hfq and Crc on an 18-mer element from the 5’-UTR of the *amiE* transcript (Pei *et al*., 2019), we hypothesized that higher order assemblies may form on longer mRNA segments that contain multiple ARN motifs in the TIR. Here, we studied Hfq/Crc assembly on mRNA fragments derived from the TIRs of the *amiE, estA* and *rbsB* genes, all of which were previously shown to be regulated by Hfq and Crc (Sonnleitner & Bläsi, 2014; Kambara *et al*., 2018). As shown in Supplementary Figure 1A and Supplementary Figure 4A, the *amiE*_*6ARN*_ sequence present in the 5’UTR of the *amiE* gene is followed by one cluster of 3 complete ARN motifs in the immediate coding region, the *amiE*_*xARN*_ motif, and two ARN triplet motifs further downstream. These downstream ARN motifs have been implicated in Hfq binding *in vivo* by proximity crosslinking of Hfq and DNA in nascent transcripts followed by Hfq-specific chromatin immunoprecipitation coupled with DNA sequencing (ChIP-seq) (Kambara *et al*., 2018). In contrast to *amiE* RNA, clusters of ARN motifs are predominantly found in the 5’-UTR of *rbsB* and *estA* mRNA, rather than in the immediate 5’coding region, which is in accord with the reported ChlP-seq data (Kambara *et al*., 2018). To verify that higher order assemblies form on different mRNA targets *in vitro*, electrophoretic mobility shift assays were performed with the mRNA fragments *amiE*_*105*_ (nts -45 to +60), *rbsB*_*110*_ (nts -75 to +33) and *estA*_*118*_ (nts -85 to +33) (Supplementary Figure 1 B). For all three transcripts, a number of higher order species was observed to form gradually with increasing Hfq concentrations (Supplementary Figure 1B, lanes 1-5). In the presence of excess Crc, formation of higher order species plateaued at around three Hfq hexamers per RNA target for all mRNA fragments tested (Supplementary Figure 1B; lanes 7-9). These observations supported the hypothesis that multiple Hfq hexamers can engage different ARN-rich motifs in a mRNA target and form defined species in the presence of Crc.

### Sequential formation of higher-order Hfq-Crc assemblies from a core complex

To characterize the details of the oligomeric state and quaternary structure, the Hfq-Crc complex formed on *amiE*_*105*_ was analyzed by cryo-EM. For the grid preparation a Hfq-titration series was performed like that used for the EMSAs shown in Supplementary Figure 1B. Figure 2A shows a gallery of the key species observed in the titration series. Stable intermediates can be resolved on the grid, and these have been ordered in the gallery in a proposed assembly pathway. Although it is possible that the pathway for the formation of the higher order complexes may be heterogenous, based on the observed species we propose a pathway where in the first step, a Hfq-Crc-Crc core is formed, whereby the *amiE*_*6ARN*_ region is bound by the Hfq distal side with two Crc molecules recognizing and engaging the Hfq-RNA complex (Figure 2A, left, 2:1:1 Hfq:*amiE*_105_:Crc). From this core, another Hfq and Crc can bind to form a higher order intermediate (Figure 2A ,middle, 2:1:3 Hfq:*amiE*_105_:Crc), and in a final step, a third Hfq engages the intermediate assembly, together with a fourth Crc molecule (Figure 2A, right, 3:1:4 Hfq:*amiE*_105_:Crc). The structure of the 3:1:4 Hfq:*amiE*_105_:Crc complex was solved at 3.6 Å resolution (GS-FSC, gold standard Fourier shell correlation (Supplementary Figure 2A, Supplementary Figure 3A-C). This complex is proposed to fully mask the a*miE* 5’-end to prevent translational initiation.

**Figure 2:**
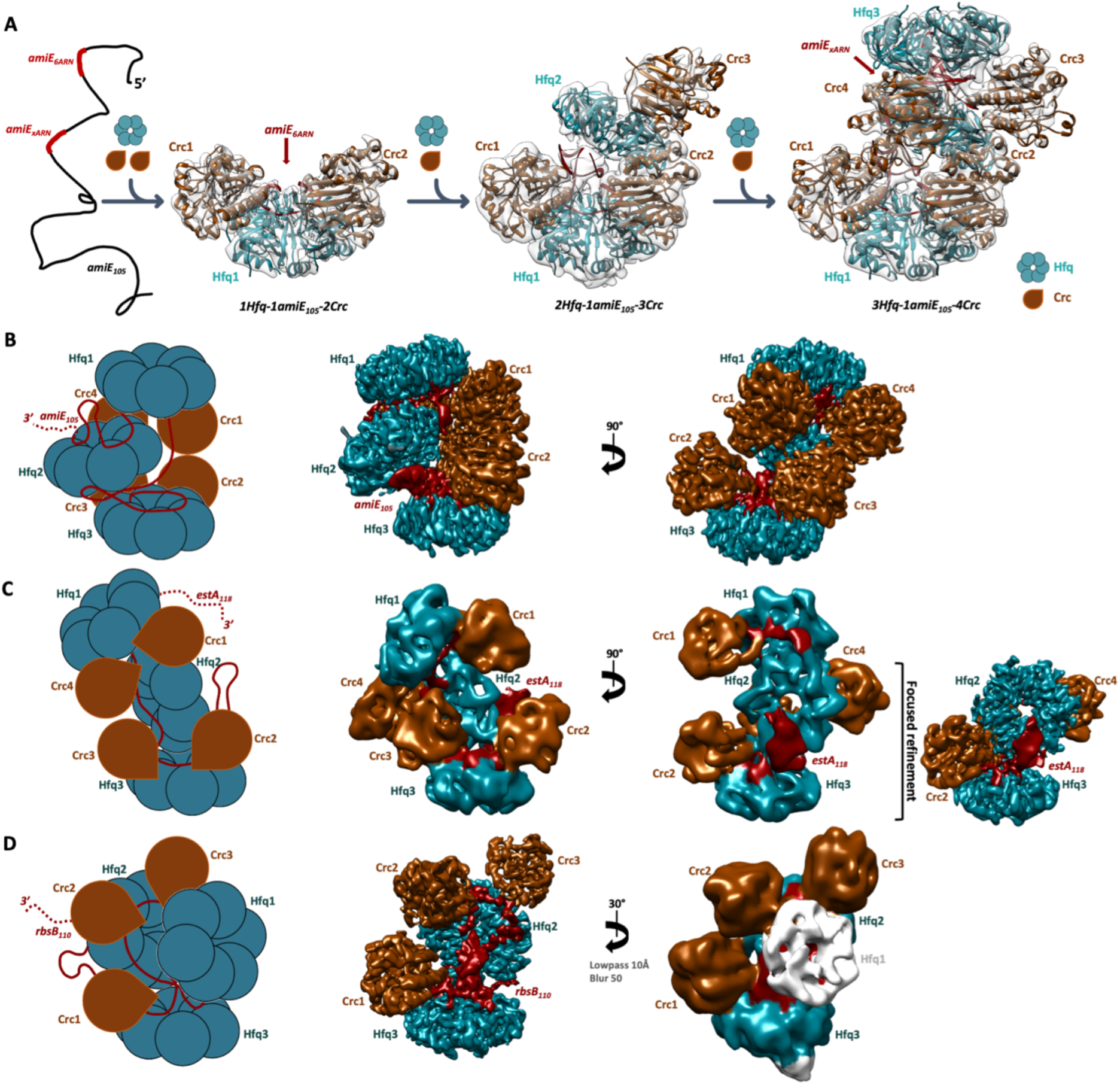
Hfq-Crc translation repression assembly pathway, and complexes formed on amiE_105_, estA_118_ and rbsB_110_. **(A)** A proposed cooperative assembly pathway for Hfq-Crc assemblies on amiE_105_. The amiE_6ARN_ region (Figure 1A) is bound by the Hfq distal side. Subsequently, two Crc molecules recognize and engage the Hfq-RNA complex (left). An additional Hfq and Crc can then bind to form a higher order intermediate (middle). In a final step, a third Hfq engages the intermediate assembly, together with a fourth Crc molecule (right). The final complex fully masks the amiE_105_ 5’-end and is proposed to inhibit translation initiation. **(B)** Schematic of the amiE_105_ translation repression complex where two Hfq hexamers (Hfq1 & 3) enclose four Crc molecules (Crc1-4) and a third Hfq hexamer (Hfq2). The amiE_105_ fragment threads through the complex, engaged by all three Hfq hexamers and all four Crc molecules. **(C)** In the estA_118_ complex two Hfq hexamers (Hfq 1 & 3) enclose four Crc molecules (Crc1-4) and a third Hfq hexamer (Hfq2), similar to the amiE_105_ complex. The estA_118_ fragment threads through the assembly and contacts all three Hfq hexamers and all four Crc molecules. Hfq1 and Crc1 are flexibly tethered to the translation repression complex and were excluded during local refinements (right panel) (see Supplementary Figure 3). Global Hfq-Crc-estA_118_ maps were lowpass filtered to 9Å to aid visualisation (middle two maps). The locally refined map is shown at 4.1 Å (right). **(D)** Three Hfq hexamers (Hfq1-3) present the rbsB_110_ mRNA to three Crc molecules (Crc1-3). The density for Hfq1 is diffuse, which may be explained by flexible association of Hfq to the translation repression complex (see Supplementary Figure 3).

### Hfq-Crc complexes are polymorphic in quaternary structure

To further elucidate the architectural principles of Hfq/Crc repressive complexes, we solved structures of Hfq-Crc complexes formed on the TIRs of the *estA*_118_ and *rbsB*_110_ mRNA segments (Supplementary Figure 1A). Representative images and 2D class averages are shown in the Supplementary Figures 2B and C, and 3D. The reconstructions were generated at 4.4 Å and 3.8 Å resolution, respectively (Figure 2C and D, Supplementary Figure 3D-I). Focused refinements resulted in a 4.1 Å resolution reconstruction of a sub-assembly of the Hfq:*estA*_*118*_:Crc complex (Figure 2C; right inset, Supplementary Figure 3E). In both complexes, the mRNA target sequences are bound by three Hfq hexamers, and three (*rbsB*_*110*_) or four (*estA*_*118*_) Crc molecules (Figure 2C-D). Notably, we observed different quaternary structures for each mRNA target, *amiE*_*105*_, *estA*_*118*_ and *rbsB*_*110*,_ demonstrating the assembly of polymorphic Hfq-Crc RNPs, driven by the mRNA sequence (Figure 2B, C and D and Supplementary Figure 4). From these structures, recurring features can be observed that define architectural principles of Hfq/Crc assembly during CCR.

In all three Hfq/Crc complexes with *amiE*_105_, *estA*_118_ and *rbsB*_110_, the ARN-rich motifs in the mRNA target sequences are bound by Hfq as described in our recent study on the Hfq-Crc-*amiE*_*6ARN*_ assembly (Pei *et al*., 2019), where the Hfq distal surface engages the ARN motifs as noted by Link *et al*. (2009). In this interaction the A- and R-site bases are bound in basic pockets on the Hfq distal side, while the N-bases are exposed, as such presenting the RNA for interaction with Crc partner molecules. Interestingly, all three complexes constitute a modular assembly that consists of one (*estA*_*118*_ *and rbsB*_*110*_) or two (*amiE*_*105*_) copies of the same Hfq-Crc-Crc core unit (Figure 3A, B and C), where a basic patch on the two Crc molecules engages mainly the phosphate backbone of the presented ARN motif (Figure 3C and D). To test the importance of this basic patch *in vivo*, translational *lacZ* reporter genes were designed for each target mRNA. The translation repression of these reporter constructs was then assessed in the presence of different Crc mutant proteins. Indeed, substitution of basic residues at the interface (Crc residues R140A and R140/141A, R138/139A) reduced translation repression to ΔCrc levels for all three reporter genes *amiE*+60*::lacZ;, estA*+18*::lacZ* and *rbsB*+13*::lacZ*) (Figure 3H). These *in vivo* results corroborate the importance of the RNA interactions for translational repression of mRNA targets by Crc. For the Hfq-*amiE*_105_-Crc assembly, the downstream *amiE*_*xARN*_ motif in the proximal coding region (Supplementary Figure 1A) only partially occupies the Hfq distal side yet still recruits 2 Crc molecules (Crc1 and 4) (Hfq 1 in Supplementary Figure 4A and Figure 3A and B, left panels). Similarly, incomplete ARN-rich motifs partially decorate the Hfq distal sides in the *estA*_*118*_ *and rbsB*_*110*_ complexes (Supplementary Figure 4B and C), recruiting one or two additional Crc molecules per Hfq hexamer to the assembly (Figure 3B). All Crc molecules bind to the presented ARN motifs *via* the basic patch described above. Together, the structures of the translation repression assemblies and the validation studies *in vivo* indicate the importance of the common RNA interactions by Crc.

**Figure 3:**
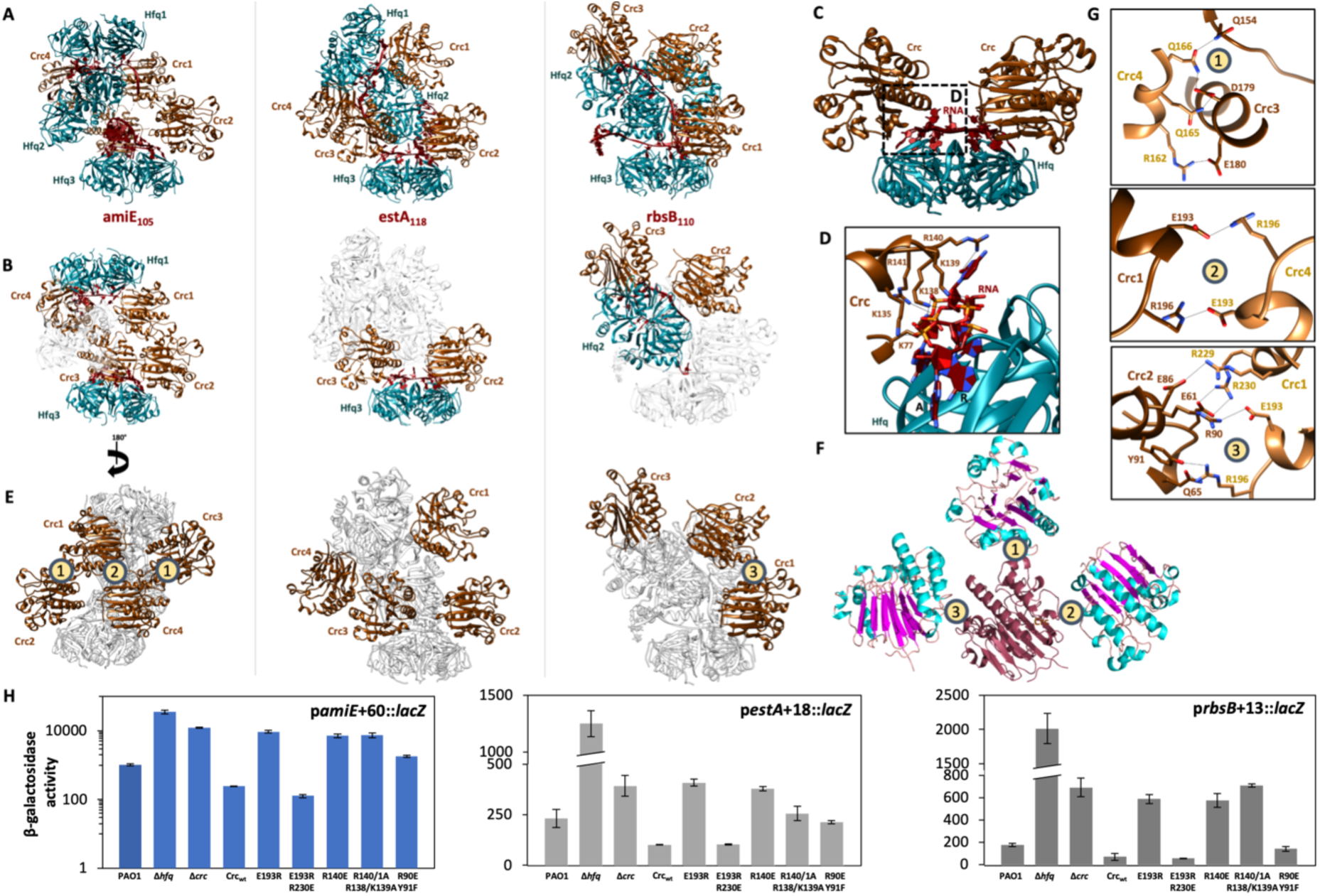
Conserved RNA binding by Crc and diverse protein-to-protein interfaces between Crc protomers in the Hfq-Crc assemblies on amiE_105_, estA_118_ and rbsB_110_. **(A-C)** All three assemblies are built from one or two Hfq-Crc-Crc core complexes, where Hfq presents – ARN-rich motifs to two Crc molecules. The full repressive assemblies formed on amiE_105_, estA_118_ and rbsB_110_ are depicted in panel A. In panel B, the same complexes are depicted, but only the Hfq-Crc-Crc core units are coloured. In panel C the common Hfq-Crc-Crc core complex is depicted. **(D)** A close-up of the Crc-RNA interactions in the core units. All Crc molecules in the repressive assemblies interact with the RNA via the same binding mode. A basic patch on the Crc surface, comprised of K77/K135/K138/K139/R140/R141, engages the phosphate backbone and a few N-site bases of the presented RNA on the Hfq distal side. **(E-G)** Crc forms homodimers in the amiE_105_ and rbsB_110_ assemblies via diverse permissive dimer interfaces. Panel E shows the full repressive assemblies formed on amiE_105_, estA_118_ and rbsB_110_ where only the Crc molecules are coloured. Three Crc dimer interfaces were observed in the amiE_105_ and rbsB_110_ complexes, annotated as 1, 2 and 3. Panel F shows the three Crc dimers observed, aligned and overlayed on a reference Crc molecule, with the interfaces annotated as in panel E. A close up of each dimer interface is given in panel G. **(H)** Translational regulation of amiE+60::lacZ, rbsB+13::lacZ and estA+18::lacZ reporter genes by Crc variants. P. aeruginosa strain PAO1Δcrc harbouring either plasmids pamiE+60::lacZ (H; left panel), pestA+18::lacZ (H, middle panel)), prbs+13::lacZ (H, right panel) or plasmid pME4510 (vector control) together with pME4510crc_Flag_ (Crc_wt_) or derivatives thereof encoding the respective mutant proteins was grown to an OD_600_ of 2.0 in BSM medium supplemented with 40 mM succinate and 40 mM acetamide (pamiE+60::lacZ) or 40 mM succinate alone (pestA+18::lacZ and prbsB+13::lacZ). The β-galactosidase values conferred by the corresponding LacZ fusion proteins are indicated. The results represent data from two independent experiments and are shown as mean and range.

In the *amiE*_*105*_ and the *rbsB*_*110*_ repressive assemblies Crc forms homodimers, and three different self-dimerization interfaces were observed (labeled as 1, 2 and 3 in Figure 3E, F, and G), constituting a second recurring feature in the repressive assemblies. Since Crc is monomeric in solution at high μM concentrations, these dimerisation interfaces must require that the Crc molecules are organized in repressive assemblies. Substitution of Glu193 for Arg is predicted to disrupt the Crc interfaces 2 and 3 in the *amiE*_105_ and *estA*_118_ assemblies due to electrostatic repulsion and impact translation repression (Figure 3G). Indeed, the E193R substitution in Crc reduced the translation repression of the *amiE*+60*::lac*Z and *rbsB*+13*::lacZ* reporter genes to ΔCrc levels (Figure 4H). Compensation of the repulsive Crc interface by the R230E substitution in Crc in turn restored translation repression for the same *amiE* and *rbsB* reporter genes (Figure 3H). Although the Crc-Crc interfaces 2 and 3 are absent in the *estA*_118_ complex, Crc residue E193 provides an alternative interaction with Hfq2 N28 (chain V) and Hfq2 R19 (chain V) that is anticipated to be weakened by the E193R substitution, which might account for the observed weakened repression effect for the *estA*+18*::lacZ* reporter gene as well (Figure 3H). The substitution R230E in Crc can form interactions with Hfq2 N28 and R19 (chain V), and can restore the Hfq/Crc interface to compensate for the E193R substitution in the *estA*_*118*_ complex. The hydrogen bonding interactions of R90 and Y91 in interface 3 of the *rbsB*_110_ complex were tested with the double mutant R90E and Y91F, and found to have roughly a 2-fold effect on translational repression (Figure 3H). Interface 3 does not occur in either the *amiE*_105_ or *estA*_118_ complexes, where instead R90 and Y91 of Crc interact with the C-terminal tail of a Hfq protomer. The double mutation R90E and Y91F de-repressed translation of the *amiE* and *estA* reporter genes roughly 7-fold and 2-fold, respectively. In summary, these results show that the Crc interaction surfaces can be directed to either form self-complementary contacts that support Crc-Crc interactions or to contact Hfq, both of which stabilize the polymorphic higher order repressive assemblies.

**Figure 4:**
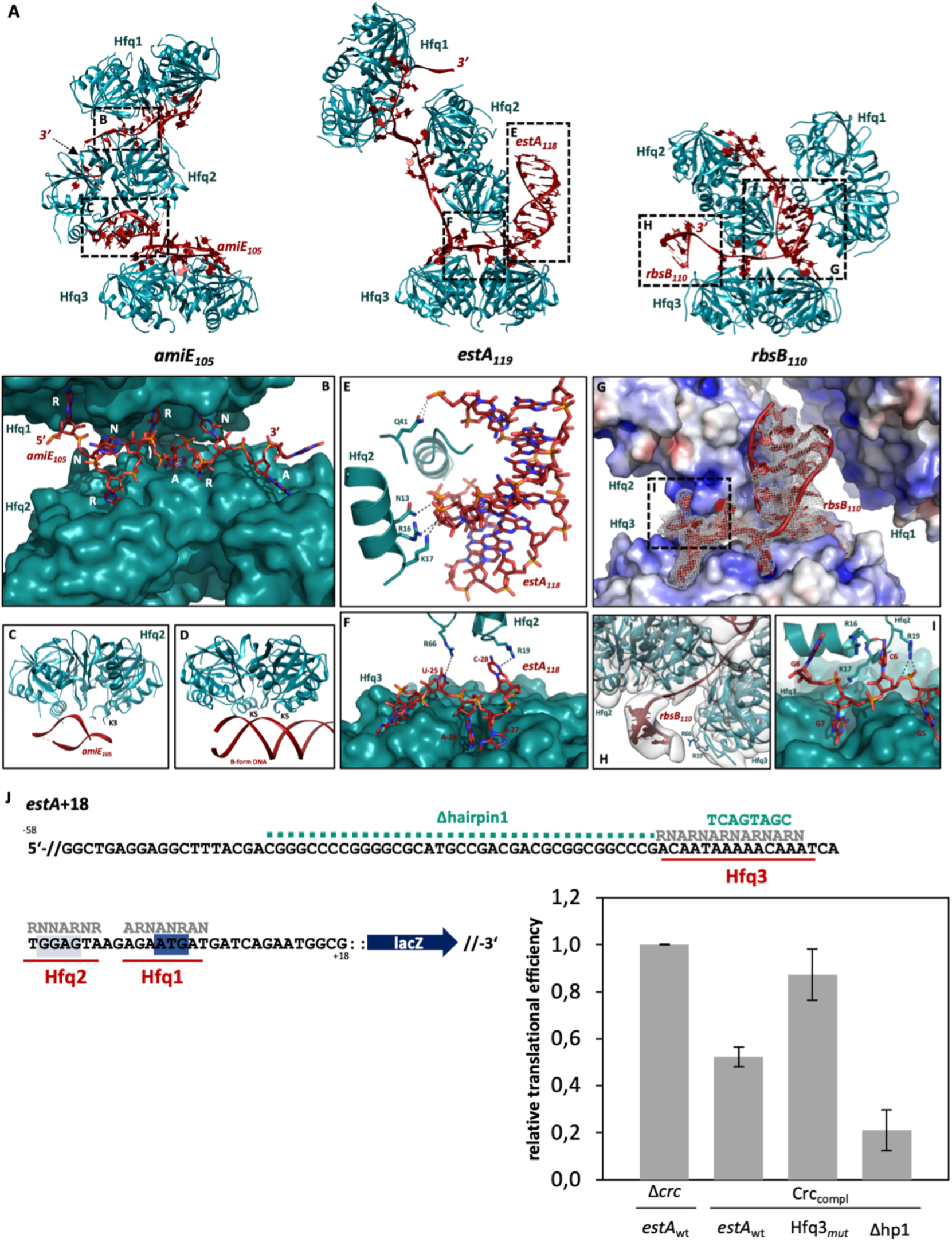
RNA sequence motifs and secondary structure elements drive oligomerization of the translation repression complexes. **(A)** Hfq-RNA components of the repressive complexes formed on amiE_105_, estA_118_ and rbsB_110_ RNAs, shown without Crc for clarity. **(B)** Close-up of the second ARN -rich motif of amiE_105_, which is shared between the Hfq1 and Hfq2 distal sides. The R-site bases in the ARN -repeats occupy alternating R-site pockets on the two Hfq distal sides, enabling higher order assembly formation. **(C)** Close-up of a hairpin structure in amiE_105_, which is coordinated by the Hfq2 proximal side (see also supplementary Figure 3A, hairpin 1). Lys3 on the proximal helix of a Hfq 2 protomer is likely to form a polar contact with the RNA backbone in the duplex. **(D)** Crystal structure of E. coli Hfq bound to B-form DNA (Orans et al., 2020), where the DNA duplex binds the proximal face of the Hfq hexamer, analogous to the amiE_105_ hairpin in (C). **(E)** Close-up of a hairpin structure in estA_105_, which is coordinated by the Hfq2 proximal side (see also Supplementary Figure 3B, hairpin1). Asn 13, Arg16 and Lys17 on the Hfq proximal helix coordinate the RNA backbone, as well as Gln41 on a proximal loop. **(F)** The ARN motif is bound to the Hfq3 distal side in the estA_119_ assembly is shared with the Hfq2 rim side. In particular, Arg19 and Arg66 form hydrogen bonds with the C-28 and U-25 bases, respectively. **(G)** A duplex-like RNA fold is coordinated by basic patches on the Hfq2 and Hfq3 distal and proximal surfaces, respectively, in the rbsB_110_ repressive complex. **(H)** At low threshold, density for a RNA duplex at the rbsB_110_ 3’-end is apparent (see also Supplementary Figure 3C hairpin1), and is coordinated by the Hfq2 proximal side. The Hfq3 rim side can make putative hydrogen bonds with the hairpin backbone via Arg66 and Arg19. **(I)** As in **(B)** and **(F)**, a short ARN motif in the rbsB_110_ sequence is shared between the Hfq3 distal side and the Hfq2 rim/proximal side. In particular, Hfq2 Arg16, Lys17 and Arg19 form contacts with the presented RNA backbone and bases in rbsB_110_. In all three complexes these Hfq-RNA-Hfq contacts drive higher order assembly formation. **(J)** Translational regulation of the estA+18::lacZ reporter gene and mutants thereof by Hfq and Crc. The mRNA sequence of the 5’UTR and the proximal coding region of estA+18::lacZ. ARN motifs bound to Hfq distal sides are annotated. The Shine Dalgarno sequence is highlighted in green and the AUG start codon is highlighted in blue. Mutations in the RNA sequence are indicated in green. Loss of the Hfq3 (Hfq3mut) binding element diminishes the repressive effects on estA mRNA significantly, whereas removal of the hairpin structure (Δhp1) results in a slight increase in translation repression. The ß-galactosidase activities were normalized to the respective mRNA levels (= relative translational efficiency) to account for intrinsic changes in mRNA stabilities eventually arising from the mutations introduced into the estA segment.

A third salient feature of the complexes is how the RNA is shared between adjacent Hfq molecules, where some Hfq distal sides present ARN motifs to the distal face or the rim of a neighbouring Hfq hexamer, rather than to Crc partner molecules. In the Hfq-*amiE*_*105*_-Crc assembly, the second *amiE*_*xARN*_ motif in the *amiE* coding region is partially shared between the distal faces of Hfq1 and Hfq2, with nucleobases occupying alternating R-site pockets on both distal sides (Figure 4A and B, Supplementary Figure 4A). This sharing of *amiE*_*xARN*_ in the *amiE* coding region between Hfq molecules drives higher-order assembly formation and efficiently masks the ribosome binding site, rationalizing the observation that the downstream ARN cluster enhances translational repression of *amiE in vivo* (Supplementary Figure 1C and D). In the Hfq-*estA*_*118*_-Crc complex, Hfq3 presents the first longer 5’ ARN motif (Supplementary Figure 4B, 12-mer, four ARN triplet repeats) to the Hfq2 rim side (Figure 4F). The Hfq2 rim-side residues Arg19 and Arg66 form hydrogen bonds with the exposed N-site bases C-28 and U-25 of *estA*_*118*_ (counting backward from the AUG start codon, with A annotated as nucleotide 1). Indeed, disruption of the ARN pattern of this Hfq3 binding site in the *estA* sequence resulted in a decrease in translation repression of the corresponding reporter gene by an order of magnitude *in vivo* (Figure 4J, Hfq3_mut_). Such disruption would also abrogate binding of *estA* by the Hfq3 distal side. A similar yet more extensive Hfq-to-Hfq presentation of the RNA target is found in the Hfq-*rbsB*_*110*_-Crc assembly, where a short RNRN-motif in the *rbsB*_*110*_ coding region is presented by the Hfq3 distal side to the Hfq2 rim/proximal side (Figure 4I). In particular, Hfq2 residues Arg16, Lys17 and Arg19 form hydrogen bonds with the phosphate backbone and the C6 nucleobase of *rbsB*_*110*_ (counting from the AUG start codon, with A annotated as nucleotide 1). From these observations, it is apparent that completion of higher-order assembly enhances translational repression, and that RNA-mediated Hfq-Hfq oligomerization drives this process.

Finally, in all three complexes RNA duplex elements interact with the Hfq proximal sides in a sequence independent manner (Figure 4A, C, E, G and H, Supplementary Figure 4). Although the local resolutions were not sufficient to resolve side chains, it is apparent that these interactions are between basic and polar residues in the proximal Hfq α-helix and the phosphate backbone of the RNA duplex structures (Figure 4C and E). In particular, the Hfq residues Lys3, Asn13, Arg16 and Lys17 are likely candidates for such interactions (Figure 4C and E). This mode of RNA secondary structure recognition by Hfq is in line with findings of a recent crystallographic study by Orans *et al*. (2020), where *E. coli* Hfq was observed to interact with the phosphate backbone of B-form DNA (Figure 4D). It is unclear what role is played by this common mode of RNA duplex-binding by Hfq in the repressive assemblies. For *estA*_118_, for example, removing the hairpin structure from the RNA construct resulted in a significantly stronger translation repression of the *estA+18::lacZ* reporter gene in *vivo*. Thus, the structural elements might confer either stabilizing or destabilizing contributions, depending on context.

In summary, the models of the repressive complexes formed on the *amiE*_105_, *estA*_*118*_ and *rbsB*_110_ transcripts and the *in vivo* reporter gene assays support a model in which the sharing of the target RNA between Hfq/Crc molecules drives formation of higher order assemblies that fully mask the ribosome binding sites. Ultimately the RNA sequence determines the quaternary structure of such translation repression complex, through blocks of ARN-repeats that can have different spacing and local imperfections (Supplementary Figure 4), and through secondary structure elements. The polymorphic character of the assembly in turn is accommodated by the hexameric character of Hfq, presenting a mosaic of basic patches on its surface, and consolidating interactions mediated through the permissive dimerization of organized Crc molecules. The different possible translation-repression assemblies, however, are all likely to fold following the recurring architectural principles described above.

## Discussion

In *P. aeruginosa* carbon catabolite repression (CCR) controls not only carbon metabolism but also other complex behaviour including infection, virulence, biofilm formation, and quorum sensing. We have shown in this study that a key component of CCR, namely the RNA chaperone Hfq, can form higher order assemblies on target mRNAs in conjunction with the protein Crc, and that such assemblies repress translation. Our cryoEM analyses reveal distinct quaternary organisations of the assemblies on the regulatory regions of the *amiE, estA* and *rbsB* mRNAs, which encode metabolic and virulence machinery and are known to be down-regulated during carbon catabolite repression (Sonnleitner & Bläsi, 2014; Kambara *et al*., 2018).

The complexes are characterised by a common core sub-assembly, comprised of one Hfq hexamer which presents an ARN-rich motif to two Crc molecules (Figure 3C). Based on the cryo-EM structures, we have elaborated rules that encode the architectural principles of translation repression assemblies. These are based on four recurring features of the complexes: i) the distal face of Hfq hexamer engages ARN-rich repeats, and is the sequence specificity determining factor. Crc can then interact with these elements *via* a distinct basic patch on its surface; ii) the proximal side of Hfq binds secondary structure elements in the RNA targets; iii) higher-order folding is driven by sharing of RNA segments between Hfq protomers and is enabled by the hexameric ring organisation of Hfq, in which protomers rich in RNA binding patches provide repetitive RNA-interaction sites. Notably, the Hfq-Hfq and Hfq-Crc interfaces formed in the translation repression complexes are almost exclusively through mutual interactions with the RNA substrate. In other words, Hfq and Crc need to bind to a polymer (RNA) for higher order assemblies to form. Polymer-bound proteins have a higher propensity to interact with each other, as part of the entropy is already lost, enabling small surfaces to contribute. iv) Some of these small surfaces are formed between neighbouring Crc molecules in the complex. These Crc dimerization interfaces support the distinct quaternary structures, and its surfaces can switch between self-complementary association or interaction with RNA/Hfq. From these principles, a diversity of quaternary structures can be supported that can regulate numerous genes in *P. aeruginosa*, with minimal proteinogenic components required.

Hfq is highly pleiotropic and involved in many riboregulatory processes. The envisaged multi-faceted control of CCR and processes linked to it entails a hidden cost of apparently requiring high numbers of Hfq and Crc in the cell, and the question arises if levels of the chaperone available are sufficient to meet the demands of forming higher order assemblies. Given that each mRNA leads to 100-1000 translated metabolic proteins on average, translational repression is only energetically beneficial for the cell if it is effective (Lynch & Marinov, 2015). Hfq levels in *P. aeruginosa* have been calculated at ∼2200 hexamers per cell during stationary phase, and nearly equimolar Crc levels per cell (∼2300) were measured (Sonnleitner & Bläsi, 2014; Sonnleitner *et al*., 2018). One scenario is that the pool is highly dynamic, and that Hfq/Crc become available through rapid turnover of transcripts to which they are bound. Indeed, transcripts are poised for degradation after translational repression (Sonnleitner & Bläsi, 2014), most likely releasing Hfq and Crc molecules that were sequestered in the repressive assembly. Secondly, genes encoding degradative proteins involved in metabolic pathways are generally only transcribed if the respective carbon source is available. This means that only a limited number of genes need to be translationally repressed in a given nutrient environment, *i*.*e*. the genes involved in metabolism of available yet non-favourable carbon sources. Lastly, it is possible that repressive responses might be graded through partial assemblies. Thus, sub-assemblies such as the Hfq-Crc-Crc core unit, might yield meaningful translation repression even when Hfq/Crc levels are insufficient to drive higher order ribonucleoprotein folding. For instance, our *in vivo* studies indicate that the downstream *amiE*_*xARN*_ -motif of *amiE* present in the coding region ensures a more complete repression, but absence of this secondary motif still results in significant translation repression (Supplementary Figure 1D).

The biological impact of these assemblies might depend on windows of opportunity arising during the synthesis of the transcript. Sequential assembly of the Hfq-Crc complexes is envisaged to occur on nascent transcripts as they emerge from the RNA polymerase, potentially coupled with RNA folding in analogy with other systems (Kambara *et al*., 2018; Yu et al. 2021). Co-transcriptional folding of the RNA, for instance the formation of stem-loop structures that influence the quaternary architectures, could affect the rates of assembly and stabilities of the complexes. Single molecule studies have shown that target RNA secondary structure presents a kinetic energy barrier that determines whether target recognition occurs before stable pairing to sRNA (Malecka and Woodson, 2021). Extending this principle, we envisage that critical kinetic steps also occur in the assembly of Hfq-Crc complexes on nascent transcripts. Indeed, removing the prominent hairpin structure that is recognised by the central Hfq in the *estA*_118_ repressive complex (Figure 4A and E, Hfq2) promotes translational repression *in vivo*, suggesting a somewhat antagonistic role for such hairpin structures in the assembly pathway. Another indication that assembly of repressive complexes might be coupled to the transcription machinery is the observation that the 3’-end of transcripts that are repressed during CCR are diminished up to 10-fold in RNA sequencing analyses (Sonnleitner et al., 2018). One mechanism that could explain this observation is the recruitment of transcription termination factors during formation of the Hfq/Crc complexes, coupling translation-repression of a transcript to termination of its transcription. This hypothesis awaits validation.

The RIL-seq results described by Kambara *et al*. (2018) and our on-grid Hfq-Crc-*amiE* assembly pathway (Figure 2A) point towards a stepwise assembly scenario, where protein binding occurs as soon as the ARN motifs are synthesized during transcription. However, given the high affinity of Hfq for RNA and the architectural principles described above, it is plausible that dynamic sampling of the RNA sequence and secondary structure elements by Hfq and/or Crc, and synergistic recruitment of multiple copies of these, forms the basis for RNP assembly. As such, co-transcriptional Hfq-Crc assembly on an RNA target might bear analogy to the synergistic co-transcriptional assembly of ribosomes on rRNA (Rodgers & Woodson 2019). If so, stepwise Hfq-Crc assembly is not sequential, but rather depends on the contextual state of the RNA binding sites and the presence of other copies of Hfq/Crc proteins. The difference here is that Hfq/Crq assembly is tuned temporally and the resulting translation repression complexes are transient in nature, subject to kinetic competition, as distinct from folding equilibrium complexes such as ribosomes and spliceosome components (Herzel *et al*., 2017, Rodgers & Woodson 2019).

The response to environmental changes and stress, and the re-routing of metabolic pathways demands systems of hierarchical control that form highly inter-connected networks. Such an intricate system is a demanding process for the cell, would require many specificity factors, *i*.*e*. protein components, to function properly. Here, we observe that specificity can be achieved with only two multifaceted protein factors and patterns in the RNA sequence. The modular nature of the protein factors and their mode of RNA interaction enables different quaternary organisations to result, *i*.*e*. less stringent folding-requirements for assembly. The regulatory outcome then becomes dependent on competitive kinetics of Hfq-Crc assembly versus translational initiation, *i*.*e*. binding of the 30S small ribosomal subunit to the mRNA RBS. Such an economic design also permits rapid modifications in network organization in the course of evolution through changes in the patterns of target RNA elements. As such, polymorphism of folded ribonucleoprotein complexes allows for a simple, highly modular regulon that underpins the noted behavioural complexity of *P. aeruginosa*.

---

**Supplementary Figure 1:**
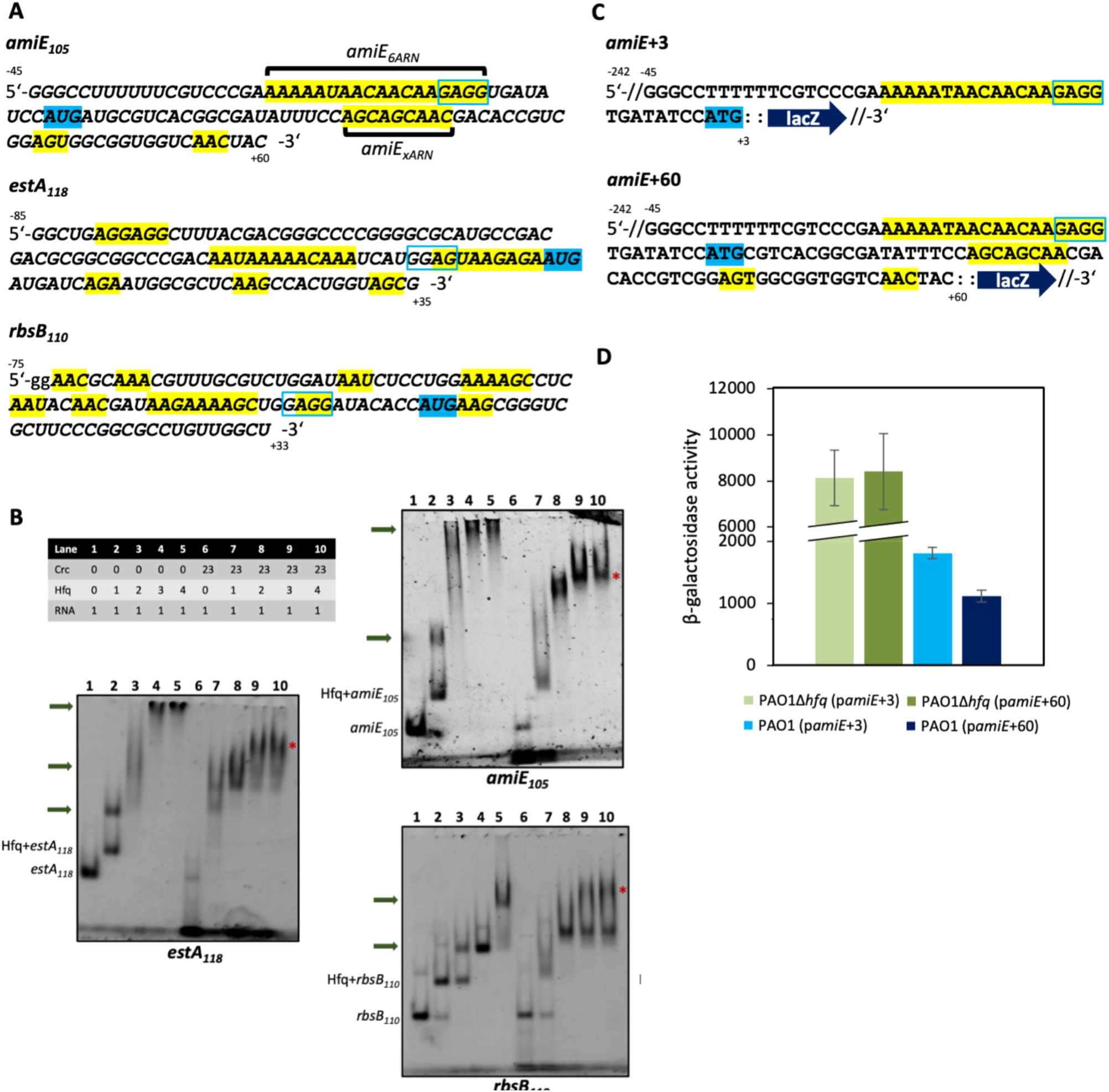
Higher order Hfq/Crc assemblies form on amiE_105_, rbsB_110_ and estA_118_ mRNA segments. **(A)** Sequences of the 5’UTRs and the proximal coding regions of amiE_105_, rbsB_110_ and estA_118_ mRNA fragments. ARN motifs are highlighted in yellow, Shine-Dalgarno sequences in blue rectangles and the AUG start codons are highlighted in blue. The amiE_6ARN_ segment used by Pei et al. (2019) is annotated as well as the shorter, secondary ARN_xARN_ segment in the amiE coding region. **(B)** Electrophoretic mobility shift assays (EMSA) with amiE_105_, estA_118_ and rbsB_110_ transcripts in the presence of Hfq and Crc. Hfq and Crc can form higher order assemblies on all RNA targets tested. Hfq molecules can engage a single RNA molecule, both in the presence and absence of Crc. Top: The table shows relative stoichiometries in the samples (RNA is at 200 nM). The protein components were mixed first, after which the RNA fragments were added. Green arrows highlight Hfq-RNA oligomers, * highlights the highest order complexes for the stoichiometry range tested. (C-D) The downstream ARN cluster enhances translational repression of amiE in vivo. **(C)** Two translational amiE::lacZ reporter genes were constructed. The amiE+3::lacZ reporter gene contained a 29-nucleotide long amiE fragment encompassing the 5’UTR with the six ARN motifs (amiE_6ARN_) and the AUG translation initiation codon fused to lacZ. The longer amiE+60::lacZ construct included additional 57 nucleotides downstream of the ATG start codon that encompass the second ARN-rich motif. **(D)** When compared with the expression in a P. aeruginosa strain lacking Hfq (PAO1Δhfq), the translation of the amiE+60::lacZ gene in the PAO1 wt strain was repressed significantly stronger than that of the amiE+3::lacZ gene. These in vivo results support the structural data presented in Figure 4A and B in that the ARN-rich motif in the coding region of amiE can contribute to the formation of effective Hfq-Crc repressive complexes in addition to the proximal amiE_6ARN_ motif.

**Supplementary Figure 2.**
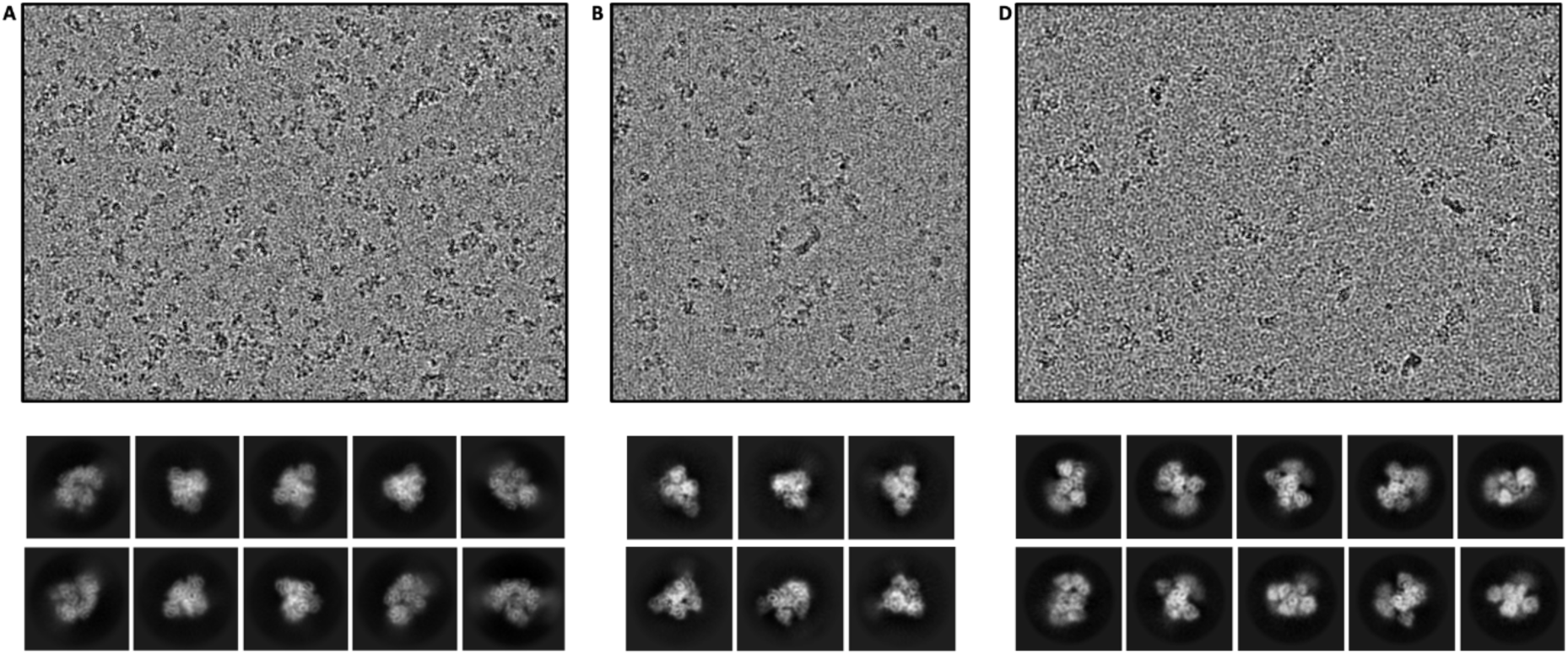
Raw images and 2D class averages of the Hfq-Crc translation repression complexes formed on amiE_105_ (A), estA_118_ (B) and rbsB_110_ (C).

**Supplementary Figure 3.**
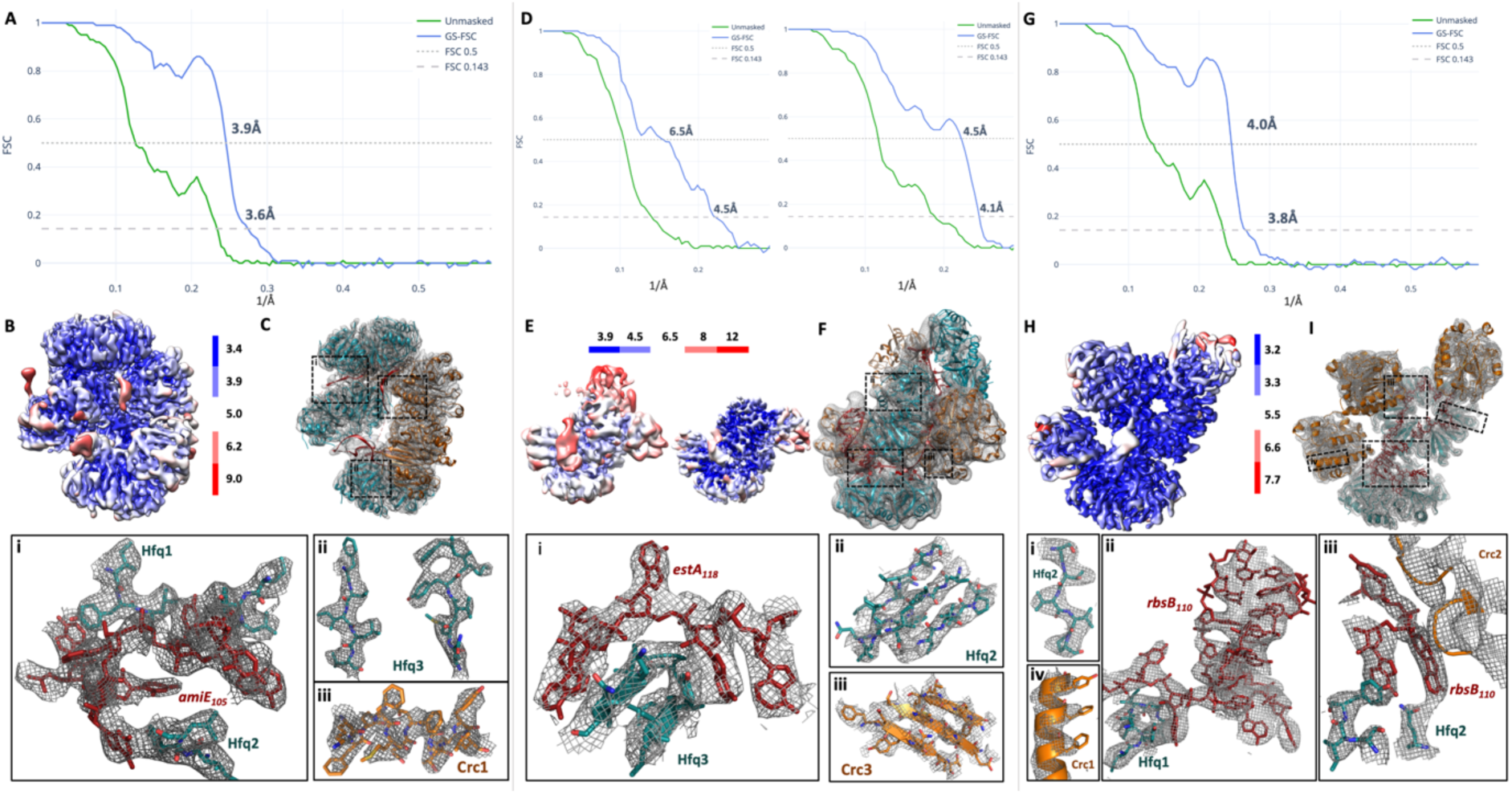
Global and local resolution analyses of the Hfq-Crc translation repression complexes formed on amiE_105_, estA_118_ and rbsB_110_. **(A), (D)**, and **(G)** show global FSC curves (gold standard) for the amiE_105_, estA_119_ and rbsB_110_ complexes, respectively. The left FSC curve in panel (D) corresponds to the global, consensus refinement, the right FSC curve corresponds to the focused refinement for the Hfq-estA119-Crc reconstruction. **(B), (E)** and **(H)** display local resolution estimates as measured by cryoSPARC at FSC 0.5.**(C)**, **(F)** and **(I)** show the respective structures, coloured as before, docked into the experimental cryo-EM maps, with the insets showing close-up of selected areas for each.

**Supplementary Figure 4:**
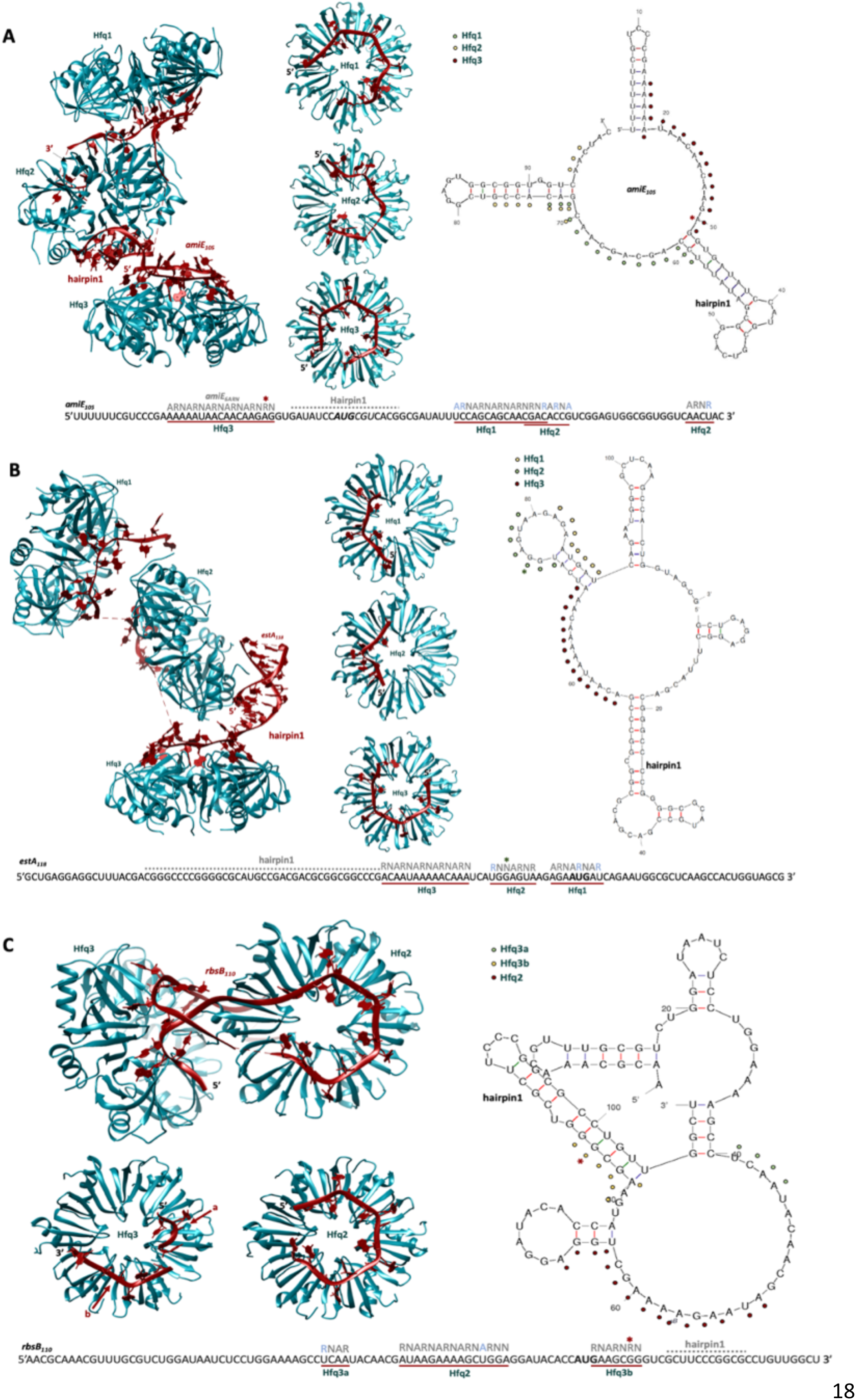
Recognition and presentation of amiE_105_, estA_119_ and rbsB_110_ by Hfq. **(A)** amiE_105_ is presented by three Hfq hexamers and adopts complete or partial ARN motif engagement on each distal side. The proximal side of Hfq2 coordinates an amiE_105_ hairpin-loop structure (hairpin1). A second hairpin-loop forms at the 3’ end of amiE_105_ on the Hfq2 distal side (hairpin2, not shown due to limited resolution). Right: annotated secondary structure prediction of amiE_105_ (mfold; Zuker, 2003). Coloured dots indicate which Hfq distal side presents the ARN-rich motif in the Hfq-amiE_105_-Crc model. An annotated sequence is depicted at the bottom of the panel. Sequences that were mapped in the cryo-EM reconstruction are underlined in red and the Hfq distal sides they bind to are labelled in green. Occupied A-, R-or N-sites are annotated in grey above each modelled sequence. * refers to an A-site ‘skipping’-violation, where the A-site on the Hfq distal site is not occupied by a base, i.e. skipped. Light blue letters refer to ‘mismatch’-violations of the ARN-rule, where a pyrimidine base occupies an A-site or R-site pocket on the Hfq distal face. The ranges for hairpin1 and hairpin2 are arbitrary due to limited local resolution in the corresponding map regions. **(B)** estA_118_ is presented by three Hfq hexamers and adopts partial ARN motif engagement on each Hfq distal side. The proximal side of Hfq2 coordinates an estA_118_ hairpin-loop structure (hairpin 1). Right: annotated secondary structure prediction of estA_118_ (mfold; Zuker, 2003). Coloured dots indicate which Hfq distal side presents the ARN-rich motif in the Hfq-Crc-estA_118_ model. An annotated sequence is depicted at the bottom of the panel. Sequences that were mapped in the cryo-EM reconstruction are underlined in red and the Hfq distal sides they bind to are labelled in green. Occupied A-, R- or N-sites are annotated in grey above each modelled sequence. * refers to an A-site ‘skipping’-violation, where the A-site on the Hfq distal site is not occupied by a base, i.e. skipped. Light blue letters refer to ‘mismatch’-violations of the ARN-rule, where a pyrimidine base occupies an A-site or R-site pocket on the Hfq distal face. **(C)** rbsB_110_ is presented by three Hfq hexamers (only the two that were well resolved in the cryo-EM maps are shown) and adopts partial ARN motif engagement on each Hfq distal side. The proximal side of Hfq2 coordinates a rbsB_110_ hairpin-loop structure (hairpin1, in the back of the Hfq2 hexamer, not annotated in the figure. Right: annotated secondary structure prediction of rbsB_110_ (mfold; Zuker, 2003). Coloured dots indicate which Hfq distal side presents the ARN-rich motif in the Hfq-rbsB_110_-Crc model. An annotated sequence is depicted at the bottom of the panel. Sequences that were mapped in the cryo-EM reconstruction are underlined in red and the Hfq distal sides they bind to are labelled in green. Occupied A-, R- or N-sites are annotated in grey above each modelled sequence. * refers to an A-site ‘skipping’-violation, where the A-site on the Hfq distal site is not occupied by a base, i.e. skipped. Light blue letters refer to ‘mismatch’-violations of the ARN-rule, where a pyrimidine base occupies an A-site or R-site pocket on the Hfq distal face. The range for hairpin1 is arbitrary due to limited local resolution in the corresponding map region.

**Table S1:**
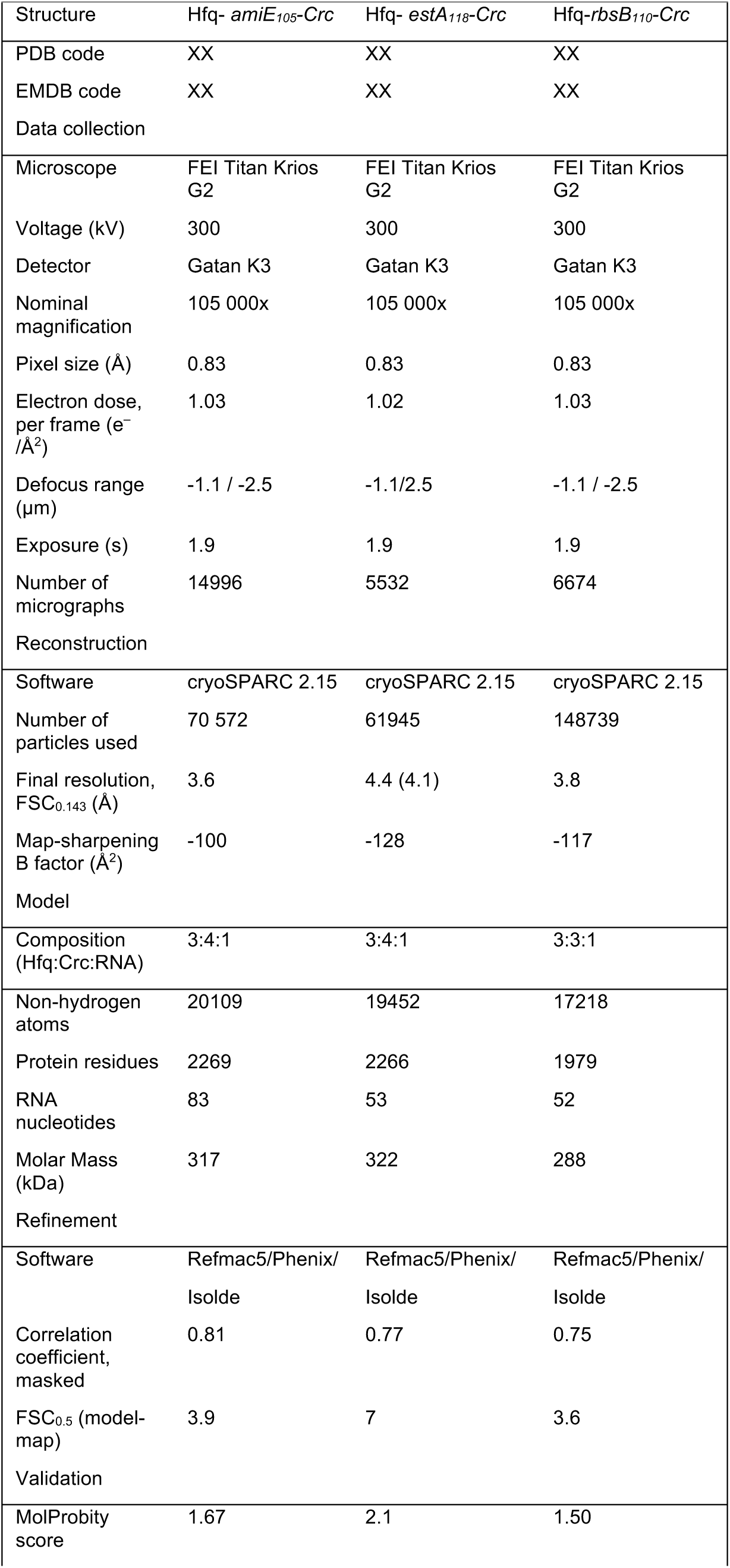
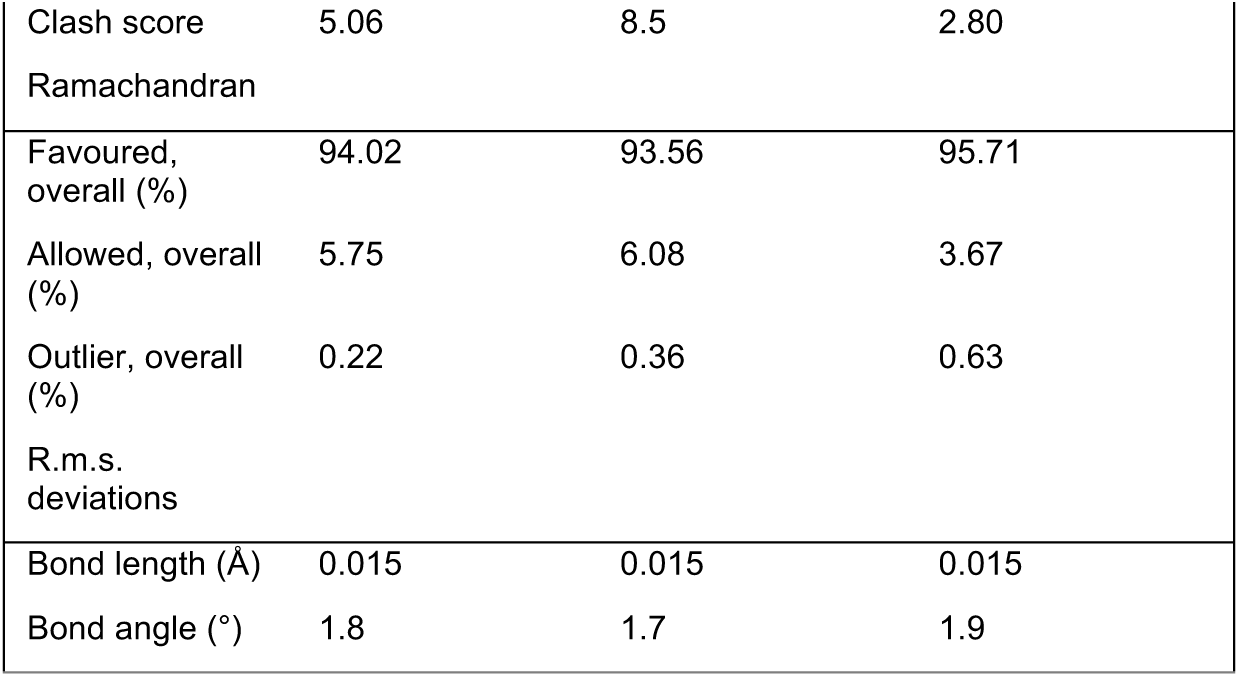
Cryo-EM data collection and refinement statistics for the Hfq/Crc/RNA structures.

## MATERIALS AND METHODS

### Purification of P. aeruginosa Hfq and Crc proteins

The purification of *P. aeruginosa* Hfq and Crc is outlined in detail in Supplementary materials.

### Bacterial strains and plasmids

The bacterial strains and plasmids used in this study are described in Supplementary materials.

### Preparation and purification of amiE_105_, rbsB_110_ and estA_118_ RNA fragments

The RNA fragments *amiE*_105_ (comprises nt -45 to +60 of *amiE* mRNA with respect to the A (+1) of the start codon), *rbsB*_110_ (comprises nt -75 to +33 of rbsB mRNA with respect to the A (+1) of the start codon plus two additional G-nucleotides at the 5’end) and *estA*_118_ (comprises nt -85 to +33 of estA mRNA with respect to the A (+1) of the start codon) were prepared by *in vitro* transcription using T7 RNA polymerase. The DNA templates were amplified by PCR using the oligonucleotide pairs R185 (5’-TCTAGACG<u>TAATACGACTCACTATA</u>GGGCCTTTTTTCGTCCCGAAAAAATAACAAC-3’) and Z172 (5’-GTAGTTGACCACCGCCACTC-3’) (*amiE*_105_), P163 (5’-AGA<u>TAATACGACTCACTATAGG</u>AACGCAAACGTTTGCGTCTGGATAATCTCCT-3’) and Q163 (5’ AGCCAACAGGCGCCGGGAAGCGACCCGCTTCAT -3’) (*rbsB*_110_) and T163 (5’-AGA<u>TAATACGACTCACTATAG</u>GCTGAGGAGGCTTTACGACGGGCCCCGGGG-3’) and U163 (5’-cgctaccagtggcttgagcgccattctgatCAT-3’) (*estA*_118_). The corresponding forward primers R185, P163 and T163, respectively, contained the T7 promoter sequence (underlined). After *in vitro* transcription with T7 RNA polymerase, the RNA fragments were gel purified using 8% polyacrylamide-8M urea gels.

### Electrophoretic mobility shift assays (EMSA)

For the EMSAs, 4 μM stocks of Hfq and Crc were prepared in binding-buffer (20 mM Tris-HCl pH 8.0, 40 mM NaCl, 10 mM KCl, 1 mM MgCl_2_), and a 2 μM stock of the RNAs was prepared in milliQ water (RNase free). The RNAs were annealed at 50°C for 3 minutes before the proteins were added. 4% poly-acrylamide (PAA) gels were used to study complex formation (6.73 ml acrylamide:bis-acrylamide, 5 ml 10x TBE, 37.7 ml milliQ water, 500 μl 10% APS and 50 μl TEMED). Hfq was titrated into a mixture of the RNAs (*amiE*_*105*_, *rbsB*_*110*_, and *estA*_*118*_) at different ratios in the presence or absence of an excess of Crc. The RNA concentration was kept constant at 200 nM. After 15 minutes of incubation at 37°C, the samples were mixed with an equal volume of loading buffer (50% v/v glycerol, 50% v/v binding buffer, 5 mM DTT), prior to loading them onto the gel. The gels were run at 4°C in 1x TBE running buffer and stained with SYBR gold.

### Cryo-EM sample preparation

For Hfq-Crc assembly on *amiE*_*105*_, the RNA was annealed at 50°C for 3 minutes. Hfq and Crc were mixed at 1.6 μM and 0.8 - 8.8 μM, respectively, prior to addition of *amiE*_*105*_ (800 nM final concentration). After incubation on ice for 1 hour, the mixture was diluted 7-fold before loading onto grids. The Hfq-*rbsB*_*110*_*-*Crc complex was prepared following a similar procedure. The *rbsB*_*110*_ fragment was annealed at 50°C for 3 minutes. Hfq and Crc were mixed at 2.8 μM and 9 μM, respectively, after which the RNA was added at 400 nM. After incubation at room temperature for 15 minutes and on ice for 1 hour, the mixture was diluted 4-fold prior to grid preparation. The Hfq-*estA*_*118*_*-*Crc complex was prepared following the same procedure as for the rbsB_110_ assembly, but the final sample was not diluted prior to grid preparation.

### Grid preparation

Graphene oxide (GO) grids were prepared from Quantifoil R1.2/1.3 grids. A 2mg/ml graphene oxide dispersion (Sigma Aldrich) was diluted 10-fold and spun down at 300 g for 30 seconds to remove large aggregates. The dispersion was then diluted 10-fold before applying 1 μl to glow-discharged grids (0.29 mbar, 15 mA, 2 minutes, Pelco Easiglow glow discharger). After drying out, the grids were stored in a grid box for 24-48 h prior to usage. 3 μl of the sample was applied to the GO grids and after 30 s of incubation, excess sample was blotted away and frozen in liquid ethane (blot force -4 to 0, blot time 3 seconds, Vitrobot markIV (Thermo Fischer)). The grids were screened on a 200 kV Talos Arctica (FEI) (Cryo-EM facility, Department of Biochemistry, University of Cambridge) and the movies were recorded on a 300 kV Titan Krios (Thermo Fischer) with either a Falcon III (Thermo Fischer) or K3 (Gatan) direct electron detector (MRC-LMB and BioCem facility, Department of Biochemistry, University of Cambridge).

### Single particle analysis, model building and refinement

All datasets were pre-processed with Warp (Tegunov & Cramer, 2019). Particle sets were cleaned up in CryoSparc (Punjani *et al*., 2017) *via* repetitive 2D classifications and heterogeneous refinements. Further extensive classifications in 2D were used to classify different assemblies observed on the grid for each of the mRNA targets. High resolution maps were generated for the highest order assemblies with non-uniform refinement in cryoSPARC (Punjani *et al*., 2019) and global- and per particle CTF refinements (Table S1). The Hfq-2Crc-*amiE*_*105*_ (147 000 particles, Figure 2A), 2Hfq-3Crc-*amiE*_*105*_ (99 000 particles, Figure 2A) and 3Hfq-4Crc-*amiE*_*105*_ (70 000 particles) assemblies were refined to 3.2Å, 3.9Å and 3.6Å respectively. The Hfq-Crc-*estA*_*118*_ map was reconstructed 4.5Å after global refinements and 4.1Å after local, masked refinements. The Hfq-Crc-*rbsB*_*110*_ map was refined to 3.8Å.

Crystal structures for *P. aeruginosa* Crc (PDB code 4JG3) and Hfq (PDB code 1U1T) were manually docked into the EM density map as rigid bodies in Chimera (Pettersen *et al*., 2004). The amiE_105_, estA_118_ and *rbsB*_*110*_ sequences were manually built into the density using Coot (Emsley *et al*., 2010). Refmac and Phenix real-space refinement with were used to iteratively refine the multi-subunit complexes, followed by manual corrections for Ramachandran and geometric outliers in Coot and ISOLDE (Table S1) (Emsley *et al*., 2010; Murshudov *et al*., 2011; Afonine *et al*., 2012; Croll, 2018). Model quality was evaluated with MolProbity (Williams *et al*., 2018).

### In vivo expression of the translational reporter genes

The ability of Hfq, Crc and Crc mutant proteins to repress the translation of the *amiE+60*::*lacZ, rbsB+13*::*lacZ* and *estA+18*::*lacZ* reporter genes was tested in the *P. aeruginosa* strains PAO1 (Holloway et al., 1979), PAO1Δ*hfq* (Sonnleitner et al., 2017) and in PAO1Δcrc (Sonnleitner et al., 2009) bearing plasmids pME4510 (vector control; Rist and Kertesz, 1998), pME4510crc_Flag_ (encodes the *crc* wt gene; Sonnleitner *et al*., 2018) or derivatives thereof encoding the Crc mutant proteins described in the text. The strains were grown to an OD_600_ of 2.0 in BSM medium (Sonnleitner *et al*., 2009) supplemented with 40 mM succinate and 40 mM acetamide (*amiE*+60::*lacZ* fusions) or only 40 mM succinate (*rbsB*+13::*lacZ* and *estA*+18::*lacZ* fusions). The β-galactosidase activities were determined as described (Miller, 1972) using cells permeabilized with 5% toluene. The β-galactosidase units in the different experiments were derived from two independent experiments.

### Determination of the relative translational efficiencies of the different estA+18::lacZ genes

To account for possible differences in mRNA stability of the different *estA+18*::*lacZ* mRNAs (Figure 4J), the relative translational efficiencies were determined by normalizing the β-galactosidase values to the mRNA levels assessed by qRT-PCR. First, total RNA was purified by the hot phenol method (Leoni et al., 1996). The remaining DNA was digested with Turbo DNase (Thermo Fisher Scientific). 2 μg of total RNA was used for cDNA synthesis with AMV reverse Transcriptase (Promega) together with 20 pmol of oligonucleotides O135 (5’-TAGCGGCTGATGTTGAACTG-3’, binds to *lacZ*) and M37 (5’-AGTCATGAATCACTCCGTGGTA-3’; binds to 16S rRNA). 5 μl of 20-fold (*lacZ*) and 1000-fold (16S rRNA) diluted cDNA samples, respectively, were used as templates for qPCR with HOT FIREpol® EvaGreen®qPCR mix (Solis biodyne) and 5 pmol of oligonucleotides (*lacZ*: O135/N135 (5’-ACTATCCCGACCGCCTTACT-3’); 16SrRNA: M37/L37 (5’-ATCGTAGTCCGGATCGCAGT-3’)) in a 20 μl reaction. The qPCR reaction was performed in a Realplex 2 Mastercylcer (Eppendorf). The PCR efficiencies and relative expression ratios of the target genes (*estA*+18::*lacZ* and mutants thereof) in comparison to the reference gene (16S rRNA) were calculated as described in Pfaffl (2001). The relative translational efficiencies were determined by normalizing the β-galactosidase values to the mRNA levels of the corresponding fusion genes and setting the relative translational efficiency of *estA*+18::*lacZ* in the absence of Crc to 1.

## Acknowledgements

The work was supported by a Wellcome Trust Investigator award to BFL (200873/Z/16/Z). TD was supported by an AstraZeneca studentship. UB and ES were supported by the Austrian Science Fund (FWF) (P28711-B22). We express our gratitude to Flavia Bassani and Armin Resch for initial experiments and the provision of materials. We thank Kasia Bandyra, Ewelina Malecka-Grajek, Nancy Standart and Alexander Borodavka for helpful discussions and advice. All grids were prepared and cryo-EM data collected at the BIOCEM facility, Department of Biochemistry, University of Cambridge. We thank Dimitri Y. Chirgadze, Steven Hardwick, and Lee Cooper for assistance with data collection at the Cryo-EM Facility.

## Supplementary materials

### P. aeruginosa Hfq purification

An Hfq-deficient *Escherichia coli* strain bearing plasmid pKEHfqPae encoding *Pseudomonas aeruginosa* Hfq (Sonnleitner *et al*., 2018) was grown in 50 ml Lysogeny broth (LB) supplemented with 0.2% wt/v glucose, 15 μg/ml kanamycin, 50 μg/ml ampicillin at 37°C in an orbital shaker at 220 rpm overnight. 5 ml of the pre-culture was used to inoculate 4 L of LB (with the same supplements) at 37°C, and the culture was grown in a shaker to an OD_600nm_ of 0.6, at which time expression of the *hfq* gene was induced with 1 mM IPTG (isopropyl β-D-1-thiogalactopyranoside). After continued growth at 4 hours, the cells were harvested by centrifugation at 5,000 g for 20 minutes at 4°C, and the pellets were resuspended in 20ml lysis buffer (50mM Tris-HCl pH 8, 1.5 M NaCl, 250 mM MgCl_2_, 1mM β-mercaptoethanol, 1 mM EDTA, 1mM PMSF) and frozen in liquid nitrogen for storage at -80°C. The thawed cells were supplemented with 20 μg/ml DNase I and lysed using an Avestin Emulsiflex C5 homogeniser (5 passes, 1000 bar). The lysate was centrifuged at 35,000 g for 20 minutes at 4°C, and the supernatant was collected and heated to 85°C in a water bath for 45 minutes. The precipitate was removed by a 20,000 g spin at 4°C for 15 minutes and 1 M (NH_4_)_2_SO_4_ was gradually added to the supernatant. The precipitate was pelleted at 20,000 g for 15 minutes (4°C), then the supernatant was filtered through a 0.42 μm Sartorius Minisart syringe filter. The sample was applied to a 5 ml HiTrap Butyl HP column (GE Lifesciences) equilibrated in buffer A (50 mM Tris-HCl pH 8, 1.5 M NaCl, 1.5 M (NH_4_)_2_SO_4_, 0.5 mM β-mercaptoethanol, 0.5 mM EDTA and 0.1 mM PMSF). After loading, 10 column volumes of buffer A were used to remove contaminants, and a 0-100% linear gradient of buffer B (50 mM Tris-HCl pH 8.0, 200 mM NaCl, 0.5 mM β-mercaptoethanol, 0.5 mM EDTA and 0.1 mM PMSF (phenylmethysulfonyl fluoride)) was used to elute Hfq. The eluted protein was diluted two-fold in buffer Hep-A (50 mM Tris-HCl pH 8.0, 100 mM NaCl, 0.5 mM β-mercaptoethanol, 0.5 mM EDTA and 0.1 mM PMSF) and loaded on a 5ml HiTrap Heparin column (GE Lifesciences) equilibrated with buffer Hep-A. 10 column volumes of Buffer Hep-A where then used to wash of any remaining contaminants. Hfq was eluted with a linear 0-60% gradient of buffer Hep-B (50 mM Tris-HCl pH 8.0, 2 M NaCl). Next, the peak fractions were pooled and concentrated in an Amicon Ultra centrifugal filter unit (10 kDa cutoff) to a final volume of 500 μl. The sample was then loaded on a Superdex 200 Increase 10/300 GL (GE Lifesciences) equilibrated with Buffer SEC-A (50mM Tris-HCl pH 7.5, 200 mM NaCl, 10% v/v glycerol). The peak fractions were flash frozen and stored at -80°C. An SDS-PAGE denaturing gel was run with the peak fractions to assess purity.

### P. aeruginosa Crc purification

Plasmid pETM14lic-6His-Crc was transformed into competent *E. coli* BL21DE3 cells by standard heat shock transformation, and the cells were plated on LB-agar plates supplemented with 50 μg/ml kanamycin. A pre-culture of the cells was grown overnight in 50 ml LB medium supplemented with 50 μg/ml kanamycin. 4 × 800 ml of LB, supplemented with 50 μg/ml kanamycin, 0.2% glucose and 2 mM Mg_2_SO_4_, were inoculated with 4 ml of the pre-culture. 3 mM of IPTG (final concentration) was used to induce expression of the *crc* gene at an OD_600nm_ of 0.6. After 3 hours, the cells were harvested by centrifugation at 5,000 g for 20 minutes. The pellet was resuspended in 50 ml Ni-A buffer (50 mM Tris-HCl pH 8.0, 300 mM NaCl, 10 mM imidazole, 1 mM β-mercaptoethanol, 0.1 mM PMSF), frozen in liquid nitrogen and stored at -80°C. The thawed cells were supplemented with 20 μg/ml DNase I and 20 μg/ml RNase A and lysed using an Avestin Emulsiflex C5 homogeniser (5 passes, 1000 bar). The lysate was centrifuged for 30 minutes at 30,000 g (4°C), and the supernatant was loaded on a 5ml HiTrap chelating column charged with NiSO_4_ and equilibrated in buffer Ni-A. The column was washed with 10 column volumes of buffer Ni-W (50 mM NaH_2_PO_4_, 300 mM NaCl, 20 mM imidazole, pH 8.0) and eluted with a linear 0-60% gradient of buffer Ni-B (50 mM NaH_2_PO_4_, 300 mM NaCl, 500 mM imidazole, pH 8.0). The peak fractions were pooled and dialyzed in 50 mM Hepes pH 8, 150 mM NaCl, 1 mM β-mercaptoethanol and the concentration was measured with a NanoDrop spectrophotometer (Thermo Fisher). For each mg of protein, 20 μg PreScission Protease (Sigma Aldrich) was added to cleave the His-tag. After 2 hours of incubation at 4°C, the sample was applied to a nickel column to remove the cleaved His-tags and PreScission Protease, and the flow through was concentrated with an Amicon Ultra centrifugal filter (5 kDa cutoff). The sample was loaded on a Superdex 200 Increase 10/300 GL (GE Lifesciences) equilibrated with Buffer SEC-A (50 mM Hepes pH 8.0, 150 mM NaCl, 1 mM TCEP (tris(2-carboxyethyl) phosphine) and 10% v/v glycerol). The peak fractions were flash frozen and stored at -80°C. An SDS-PAGE denaturing gel was run with the peak fractions to assess purity.

### Bacterial strains / plasmids used in this study and construction of lacZ-reporter genes

To construct translational gene fusions between *amiE* and *lacZ*, DNA fragments containing nucleotides from -242 to +3 (*amiE*+3) and to +60 (*amiE*+60), respectively, with regard to the A (+1) of the start codon of *amiE*, were amplified by PCR using the oligonucleotides A1 (5’-TTTTTTGAATTCGGCTGCATGCTATCTCAGGCGC-3’) and either F173 (*amiE*+3; 5’-TTTTTTCTGCAGGTAGTTGACCACCGCCACTC-3’) or G173 (*amiE*+60; 5’-TTTTTTCTGCAGCATGGATATCACCTCTTGTTG-3’) and chromosomal DNA of strain PAO1 as template. The PCR fragments were cleaved with *Eco*RI and *Pst*I and then ligated into the corresponding sites of plasmid pME6015 (Schnider-Keel et al., 2000), generating plasmids p*amiE*+3::*lacZ* and p*amiE*+60::lacZ, respectively.Plasmid p*rbsB*+13::*lacZ* was constructed as described in Kambara *et al*. (2018). A DNA fragment containing nucleotides from -372 to +13 with regard to the A (+1) of the start codon of *rbsB*, was amplified by PCR using the oligonucleotide pair T191 (5’-ATATGAATTCGTCCAGCCTGGAGGTCTACAAG-3’) and U191 (5’-ATATCTGCAGCGACCCGCTTCATGGTG-3’) and chromosomal DNA of strain PAO1 as template. The PCR fragments were cleaved with *Eco*RI and *Pst*I and then ligated into the corresponding sites of plasmid pME6015 (Schnider-Keel et al., 2000), generating plasmid p*rbsB*+13::*lacZ*.

The construction of p*estA*+18::*lacZ* has been described by Sonnleitner & Bläsi (2014), wherein it was termed pTLestA. Plasmid p*estA*+18::lacZ contains a DNA fragment of *estA* spanning nucleotides -580 to +18 with regard to the A (+1) of the start codon of *estA*.

Plasmid pTLestA-ΔCA (herein termed pestA+13ΔCA::lacZ), wherein a part of the Hfq3 binding site (AAAACAA) in *estA* was mutated to TCAGTAGC (Hfq3_mut_ in Figure 4) has been described in Sonnleitner *et al*. (2012)).

Plasmid p*estA*+18-Δhp1::*lacZ* was constructed employing overlapping PCR. The PCR fragments were amplified with primer pairs Q67 (5’-TTTTTGAATTCGAGCAGCCTGGCACGC-3’)/G191 (5’-TCGTAAAGCCTCCTCAG-3’) and H191 (5’-CTGAGGAGGCTTTACGAACAATAAAAACAAATCATGGAGTAAGAGA-3’)/R67 (5’-TTTTGGATCCGAGCGCCATTCTGATCAT-3’) and p*estA*+18::*lacZ* as template. The PCR fragments were combined and used as a template for a second overlapping PCR with primers Q67 and R67. The resulting PCR fragment, comprising fragment of *estA* from nucleotide -580 to +18 with a deletion of 37 nucleotides from nucleotides -66 to -30 with regard to the A (+1) of the start codon of *estA* was digested with *Eco*RI and *Bam*HI and ligated into the corresponding sites of pME6015.

### Construction of plasmids encoding Crc variant proteins

Derivatives of plasmid pME4510crc_Flag_ (Sonnleitner *et al*., 2018) were constructed by site directed mutagenesis using plasmid pME4510crc_Flag_ as template and the mutagenic oligonucleotide pairs E191 (5’-GAGCAAGCAGCGTGCCGCGGCCGCCGAATACATCTACTGC-3’)/ F191 (5’-GCAGTAGATGTATTCGGCGGCCGCGGCACGCTGCTTGCTC-3’) (Crc_R138A,K139A,R140A,R141A_), N191 (5’-CGATCGTTACGGGGAATTCCTGCAAGCCGACTTCGACAAGG-3’)/ O191 (5’-CCTTGTCGAAGTCGGCTTGCAGGAATTCCCCGTAACGATCG-3’) (Crc_R90E,Y91F_), L191 (5’-CCTTTATGCCTGCGATGCCCGTCTACCCGAACAGG-3’)/ M191 (5’-CCTGTTCGGGTAGACGGGCATCGCAGGCATAAAGG-3’) (CrcE61R) and A191 (5’-CTTAGGTTTCCGCACGGCCGATC-3’)/ B191 (5’-GATCGGCCGTGCGGAAACCTAAG-3’) (CrcE86R), respectively. The parental plasmid templates were digested with DpnI and the mutated nicked circular strands were transformed into *E. coli* XL1-Blue, generating plasmid pME4510crc_(R138A,K139A,R140A,R141A)Flag_, pME4510crc_(R90E,Y91F)Flag_ and pME4510crc_(E61R,E86R)Flag_.

The construction of plasmids pME4510crc_(E193R)Flag_, pME4510crc_(R230E)Flag_, pME4510crc_(E193R,R230E)Flag_, pME4510crc_(E193A,R230E)Flag_, pME4510crc_(R229A,R230E)Flag_ and pME4510crc_(R140E)Flag_ as described in Pei *et al*. (2019).

## References

Abdou, L., Chou, H.T., Haas, D. and Lu, C.D. (2011) Promoter recognition and activation by the global response regulator CbrB in Pseudomonas aeruginosa. J Bacteriol, 193, 2784–2792.

Afonine, P.V., Grosse-Kunstleve, R.W., Echols, N., Headd, J.J., Moriarty, N.W., Mustyakimov, M., Terwilliger, T.C., Urzhumtsev, A., Zwart, P.H., and Adams, P.D. (2012). Towards automated crystallographic structure refinement with phenix.refine. Acta Crystallogr. D Biol. Crystallogr. 68, 352–367.

Ali Azam, T., Iwata, A., Nishimura, A., Ueda, S., & Ishihama, A. (1999). Growth phase-dependent variation in protein composition of the Escherichia coli nucleoid. Journal of bacteriology, 181(20), 6361–6370. https://doi.org/10.1128/JB.181.20.6361-6370.1999

Burnley, T., Palmer, C. M., & Winn, M. (2017). Recent developments in the CCP-EM software suite. Acta crystallographica. Section D, Structural biology, 73(Pt 6), 469–477. https://doi.org/10.1107/S2059798317007859

Corona, F., Reales-Calderon, J.A., Gil, C. and Martinez, J.L. (2018) The development of a new parameter for tracking post-transcriptional regulation allows the detailed map of the Pseudomonas aeruginosa Crc regulon. Sci Rep, 8, 16793.

Croll, T.I. (2018). ISOLDE: a physically realistic environment for model building into low-resolution electron-density maps. Acta Cryst. D74

Dubey, A.K., Baker, C.S., Romeo, T. and Babitzke, P. (2005) RNA sequence and secondary structure participate in high-affinity CsrA-RNA interaction. RNA, 11, 1579–1587.

Emsley, P., Lohkamp, B., Scott, W.G., and Cowtan, K. (2010). Features and development of Coot. Acta Crystallogr. D Biol. Crystallogr. 66, 486–501.

Fernandez, L., Breidenstein, E.B., Taylor, P.K., Bains, M., de la Fuente-Nunez, C., Fang, Y., Foster, L.J., and Hancock, R.E. (2016). Interconnection of post-transcriptional regulation: the RNA-binding protein Hfq is a novel target of the Lon protease in Pseudomonas aeruginosa. Sci. Rep. 6, 26811.

Gebhardt, M.J., Kambara, T.K., Ramsey, K.M. and Dove, S.L. (2020) Widespread targeting of nascent transcripts by RsmA in Pseudomonas aeruginosa. Proc Natl Acad Sci U S A, 117, 10520–10529.

Goodman, A.L., Merighi, M., Hyodo, M., Ventre, I., Filloux, A. and Lory, S. (2009) Direct interaction between sensor kinase proteins mediates acute and chronic disease phenotypes in a bacterial pathogen. Genes Dev, 23, 249–259.

Herzel, L., Ottoz, D.S.M., Alpert, T., and Neugebauer, K.M. (2017). Splicing and transcription touch base: co-transcriptional spliceosome assembly and function. Nat. Rev. Mol. Cell. Biol. 18, 637–650.

Holmqvist, E., Wright, P.R., Li, L., Bischler, T., Barquist, L., Reinhardt, R., Backofen, R. and Vogel, J. (2016) Global RNA recognition patterns of post-transcriptional regulators Hfq and CsrA revealed by UV crosslinking in vivo. EMBO J, 35, 991–1011.

Huang, J., Sonnleitner, E., Ren, B., Xu, Y., and Haas, D. (2012). Catabolite repression control of pyocyanin biosynthesis at an intersection of primary and secondary metabolism in Pseudomonas aeruginosa. Appl. Environ. Microbiol. 78, 5016–5020.

Jakobi, A.J., Wilmanns, M. & Sachse, C. 2017. Model-based local density sharpening of cryo-EM maps. Elife. 6, 1–26.

Kambara, T.K., Ramsey, K.M. and Dove, S.L. (2018) Pervasive targeting of nascent transcripts by Hfq. Cell Rep, 23, 1543–1552.

Krepl, M., Dendooven, T., Luisi, B. F., & Sponer, J. (2021). MD simulations reveal the basis for dynamic assembly of Hfq-RNA complexes. J. Biol. Chem, 296, 100656. https://doi.org/10.1016/j.jbc.2021.100656

Leoni, L., Ciervo, A., Orsi, N. and Visca, P. (1996) Iron-regulated transcription of the pvdA gene in Pseudomonas aeruginosa: effect of Fur and PvdS on promoter activity. J Bacteriol.178, 2299–2313.

Lu, P., Wang, Y., Zhang, Y., Hu, Y., Thompson, K.M., and Chen, S. (2016). RpoS-dependent sRNA RgsA regulates Fis and AcpP in Pseudomonas aeruginosa. Mol. Microbiol. 102, 244–259.

Lynch, M., & Marinov, G. K. (2015). The bioenergetic costs of a gene. Proceedings of the National Academy of Sciences of the United States of America, 112, 15690–15695. https://doi.org/10.1073/pnas.1514974112

Milojevic, T., Grishkovskaya, I., Sonnleitner, E., Djinovic-Carugo, K. and Bläsi, U. (2013) The Pseudomonas aeruginosa catabolite repression control protein Crc is devoid of RNA binding activity. PLoS One, 8, e64609.

Malecka, E.M. and Woodson, S.A. (2021) Stepwise sRNA targeting of structured bacterial mRNAs leads to abortive annealing. Mol. Cell 81, 1988–1999.

Malecka, E.M., Bassani, F., Dendooven, T., Sonnleitner, E., Rozner, M., Albanese, T.G., Resch, A., Luisi, B.F., Woodson, S. and Bläsi, U. (2021). Stabilization of Hfq-mediated translational repression by the co-repressor Crc in Pseudomonas aeruginosa. Nucleic Acids Res. 49, 7075–7087.

Moreno, R., Hernandez-Arranz, S., La Rosa, R., Yuste, L., Madhushani, A., Shingler, V. and Rojo, F. (2015) The Crc and Hfq proteins of Pseudomonas putida cooperate in catabolite repression and formation of ribonucleic acid complexes with specific target motifs. Environ Microbiol, 17, 105–118.

Murshudov, G.N., Skubák, P., Lebedev, A.A., Pannu, N.S., Steiner, R.A., Nicholls, R.A., Vagin, A.A. (2011). REFMAC5 for the refinement of macromolecular crystal structures. Acta Crystallogr. D Biol. Crystallogr. 67, 355–367.

O’Toole, G.A., Gibbs, K.A., Hager, P.W., Phibbs, P.V. Jr., and Kolter, R. (2000). The global carbon metabolism regulator Crc is a component of a signal transduction pathway required for biofilm development by Pseudomonas aeruginosa. J. Bacteriol. 182, 425–431.

Pei, X.Y., Dendooven, T., Sonnleitner, E., Chen, S., Bläsi, U. and Luisi, B.F. (2019) Architectural principles for Hfq/Crc-mediated regulation of gene expression. Elife, 8, e43158.

Pettersen, E.F., Goddard, T.D., Huang, C.C., Couch, G.S., Greenblatt, D.M., Meng, E.C., and Ferrin, T.E. (2004). UCSF Chimera - a visualization system for exploratory research and analysis. J. Comput. Chem. 25, 1605–1612.

Pfaffl, M.W. (2001) A new mathematical model for relative quantification in real-time RT-PCR. Nucleic Acid Res. doi: 10.1093/nar/29.9.e45.

Punjani, A., Rubinstein, J.L., Fleet, D.J., and Brubaker, M.A. (2017). cryoSPARC: algorithms for rapid unsupervised cryo-EM structure determination. Nat. Methods 14, 290–296.

Punjani A., Zhang H., Fleet D.J. (2019) Non-uniform refinement: Adaptive regularization improves single particle cryo-EM reconstruction. bioRxiv, doi:10.1101/2019.12.15.877092.

Pusic, P., Tata, M., Wolfinger, M.T., Sonnleitner, E., Häussler, S. and Bläsi, U. (2016). Cross-regulation by CrcZ RNA controls anoxic biofilm formation in Pseudomonas aeruginosa. Sci. Rep. 6, 39621.

Pusic, P., Sonnleitner, E., Krennmayr, B., Heitzinger, D.A., Wolfinger, M.T., Resch, A. and Bläsi, U. (2018) Harnessing Metabolic Regulation to Increase Hfq-Dependent Antibiotic Susceptibility in Pseudomonas aeruginosa. Front Microbiol. 9, 2709

Pusic, P., Sonnleitner, E., and Bläsi, U. (2021) Specific and global RNA regulators in Pseudomonas aeruginosa. Int. J. of Molec. Sci. doi: 10.3390/ijms22168632.

Ramírez-Aportela, E., Vilas, J.L., Melero, R., Conesa, P. et al. 2020. Automatic local resolution-based sharpening of cryo-EM maps. Bioinformtics 36, 765–772.

Rodgers, M. L., & Woodson, S. A. (2019). Transcription Increases the Cooperativity of Ribonucleoprotein Assembly. Cell 179, 1370–1381.e12. https://doi.org/10.1016/j.cell.2019.11.007

Rojo, F. (2010) Carbon catabolite repression in Pseudomonas: optimizing metabolic versatility and interactions with the environment. FEMS Microbiol Rev. 34, 658–684.

Romero, M., Silistre, H., Lovelock, L., Wright, V.J., Chan, K.G., Hong, K.W., Williams, P., Camara, M. and Heeb, S. (2018) Genome-wide mapping of the RNA targets of the Pseudomonas aeruginosa riboregulatory protein RsmN. Nucleic Acids Res. 46, 6823–6840.

Santiago-Frangos, A. and Woodson, S.A. (2018) Hfq chaperone brings speed dating to bacterial sRNA. Wiley Interdiscip. Rev. RNA. doi: 10.1002/wrna.1475.

Santiago-Frangos, A., Frohlich, K.S., Jeliazkov, J.R., Malecka, E.M., Marino, G., Gray, J.J., Luisi, B.F, Woodson, S.A. and Hardwick, S.W. (2019) Caulobacter crescentus Hfq structure reveals a conserved mechanism of RNA annealing regulation. Proc. Natl. Acad. Sci. U.S.A.. 116, 10978–10987.

Schubert, M., Lapouge, K., Duss, O., Oberstrass, F.C., Jelesarov, I., Haas, D. and Allain, F.H. (2007) Molecular basis of messenger RNA recognition by the specific bacterial repressing clamp RsmA/CsrA. Nat. Struct. Mol. Biol. 14, 807–813.

Sonnleitner, E., Hagens, S., Rosenau, F., Wilhelm, S., Habel, A., Jäger, K.E., and Bläsi, U. (2003) Reduced virulence of a hfq mutant of Pseudomonas aeruginosa O1. Microb. Pathog. 35, 217–228.

Sonnleitner, E., Pusic, P., Wolfinger, M.T. and Bläsi, U. (2020) Distinctive Regulation of carbapenem susceptibility in Pseudomonas aeruginosa by Hfq. Front. Microbiol. 11, 1001.

Sonnleitner, E., Schuster, M., Sorger-Domenigg, T., Greenberg, E.P., and Bläsi, U. (2006). Hfq-dependent alterations of the transcriptome profile and effects on quorum sensing in Pseudomonas aeruginosa. Mol. Microbiol. 59, 1542–1558.

Sonnleitner, E. and Bläsi, U. (2014) Regulation of Hfq by the RNA CrcZ in Pseudomonas aeruginosa carbon catabolite repression. PLoS Genet, 10, e1004440.

Sonnleitner, E., Wulf, A., Campagne, S., Pei, X.Y., Wolfinger, M.T., Forlani, G., Prindl, K., Abdou, L., Resch, A., Allain, F.H. et al. (2018) Interplay between the catabolite repression control protein Crc, Hfq and RNA in Hfq-dependent translational regulation in Pseudomonas aeruginosa. Nucleic Acids Res, 46, 1470–1485.

Tegunov, D., Cramer, P. (2019). Real-time cryo-electron microscopy data preprocessing with Warp. Nat. Methods 16, 1146–1152. https://doi.org/10.1038/s41592-019-0580-y

Valentini, M., Garcia-Maurino, S.M., Perez-Martinez, I., Santero, E., Canosa, I. and Lapouge, K. (2014) Hierarchical management of carbon sources is regulated similarly by the CbrA/B systems in Pseudomonas aeruginosa and Pseudomonas putida. Microbiology 160, 2243–2252.

Williams, C., Headd, J., Moriarty, N., Prisant, M., Videau, L., Deis, L., Verma, V., Keedy, D., Hintze, B., Chen, V., Jain, S., Lewis, S., Arendall, W., Snoeyink, J., Adams, P., Lovell, S., Richardson, J. and Richardson, D. (2017). MolProbity: More and better reference data for improved all-atom structure validation. Protein Science 27, 293–315.

Winsor G. L., Griffiths E. J., Lo R., Dhillon B. K., Shay J. A., Brinkman F. S. L. (2016). Enhanced annotations and features for comparing thousands of Pseudomonas genomes in the Pseudomonas genome database. Nucleic Acids Res. 44, 646–653. 10.1093/nar/gkv1227

Yang, N., Ding, S., Chen, F., Zhang, X., Xia, Y., Di, H., Cao, Q., Deng, X., Wu, M., Wong, C., Tian, X., Yang, C., Zhao, J. and Lan, L. (2015). The Crc protein participates in down-regulation of the lon gene to promote rhamnolipid production and quorum sensing in Pseudomonas aeruginosa. Mol.Microbiol. 96, 526–547.

Yu, A.M., Gasper, P.M., Cheng, L., Lai, L.B., Kaur, S., Gopalan, V., Chen, A.A. and Lucks, J.B. (2021) Computational reconstructing cotranscriptional RNA folding from experimental data reveals rearrangement of non-native folding intermediates. Mol. Cell 81, 870–883.

Zhang, K. (2016). Gctf: real-time CTF determination and correction. J. Struct. Biol. 193, 1–12.

Zhang, L., Chiang, W.C., Gao, Q., Givskov, M., Tolker-Nielsen, T., Yang, L., and Zhang, G. (2012). The catabolite repression control protein Crc plays a role in the development of antimicrobial-tolerant subpopulations in Pseudomonas aeruginosa biofilms. Microbiology 158, 3014–3019.

Zhang, Y.F., Han, K., Chandler, C.E., Tjaden, B., Ernst, R.K. and Lory, S. (2017) Probing the sRNA regulatory landscape of Pseudomonas aeruginosa: post-transcriptional control of determinants of pathogenicity and antibiotic susceptibility. Mol. Microbiol. 106, 919–937.

Zheng, S.Q, Palovcak, E., Armache, J.-P., Cheng, Y. and Agard, D.A. (2017). MotionCor2: anisotropic correction of beam-induced motion for improved cryo-electron microscopy. Nat. Methods 14, 331–332.

